# Long-read RNA-seq delineates temporal transcriptional dynamics in multiplexed and sexed single medfly embryos

**DOI:** 10.1101/2025.10.29.685472

**Authors:** Anthony Bayega, Spyros Oikonomopoulos, Dimitris Rallis, Dionisia Mavritsakis, Antonia Spanomitrou, Kostas D Mathiopoulos, Jiannis Ragoussis

## Abstract

Long-read RNA sequencing has great potential to improve genomic characterization of non-model organisms due to its ability to yield full-length genes. Coupled with absolute gene expression quantification, dynamics of development orchestrated at transcript level can be elucidated with high precision. The resolution of this precision can be further improved by studying organisms as close as possible to their basic entities, single cells for example or single embryos. Here, we collected developing embryos of the Mediterranean fruit fly (medfly, *Ceratitis capitata*) at hourly time-points for the first 15 hours of development. The medfly is an organism of huge economic importance in agriculture due to its wide host range including apples, pear, citrus, olives, etc. We simultaneously isolated total RNA and genomic DNA from single embryos and sexed the embryos using Y-specific PCR assays. The RNA, spiked with external ERCC standards to aid in absolute quantification, was used to perform Nanopore long-read RNA-seq. We developed a genome-guided transcriptome assembly based on full-length transcripts and identified a total of 22,875 transcripts comprising 3879 novel genes, missed in the current NCBI predicted gene models. We show that, indeed, the absolute quantification of gene expression performs superiorly to relative quantification in highly dynamic systems such as developing embryos. Further, we used unsupervised clustering and lineage tracing algorithms to group and accurately place embryos along a pseudo-temporal development trajectory. We show that medfly embryos undergo successive waves of zygotic genome activation. We discover a dramatic reorganization of maternally deposited mRNA occurring within the first 3 hours of egg laying followed by maternal-to-zygotic transition. We finally identify modules of temporal synexpression and elucidate the biological role of these modules. Together, these results provide the first detailed look at early embryo development in the medfly and should aid in future control efforts of this pest.

## Introduction

cDNA sequencing using short-read technologies such as sequence-by-synthesis (Illumina Inc., USA) is currently the standard in high-throughput transcriptional exploration. Due to their short-read nature however, these technologies are inefficient in delineating the full breadth of the transcriptome landscape particularly regarding isoform diversity [1, 2]. We and others have shown that long-read technologies provide full-length transcript resolution [3, 4] and enable identification of hitherto unknown genes and isoforms [5–8]. This can be particularly useful for organisms that are not well characterized at a genomics level. Among long-read technologies, the Oxford Nanopore Technologies (ONT, UK) protein nanopore sequencing is particularly attractive due to high throughput, simpler library preparation workflow, and gene expression quantification that is comparable to current standards [5–8].

Gene expression is a tightly regulated process. In highly dynamic biological systems like developing embryos, precise and coordinated dramatic shifts in transcriptome kinetics occur in quick succession. Although relative normalization is a common method of measuring and comparing transcriptional changes, direct absolute normalization has been shown to perform superiorly to relative normalization in capturing such dynamic transcriptional changes [9]. Absolute normalization can be achieved by addition of external RNA standards such as the ERCC [10]. Coupled with time-course experimentation, absolute normalization enables direct measurement of transcript kinetics thus providing a quantitative understanding of the rate of change of transcript copy numbers with time. Such a quantitative understanding of the transcriptome during embryogenesis can greatly improve our understanding of the impact of different transcript regulation strategies on gene expression. Further, there has been a recent appreciation of studying biological systems at single cell level. In insects, embryos go through a syncytial blastoderm stage where nuclei divide and multiply in a common medium prior to the cellular blastoderm stage. In such systems, studying of single embryos is ideal to elucidate biological mechanisms and also account for heterogeneity in biological systems. In previous studies, elucidation of transcriptional changes in developing embryos has relied on manual staging of embryos or collecting embryos at specific time points following egg-laying or fertilization [11–13]. However, embryos laid as a batch can be at different developmental times because of differences in actual time of fertilization or being withheld within the fly prior to being laid. Single-cell RNA-seq methods that have been developed to cluster cells based on their transcription profile can provide more accurate staging and lineage tracing [12]. Furthermore, sexing of these embryos can aid in delineating sex-specific biological mechanisms.

Among insects, the Tephritidae family contains some of the most important agricultural pests, such as the oriental fruit fly (*Bactrocera dorsalis*), the Mexican fruit fly (*Anastrepha ludens*), the Australian Q-fly (*Bactrocera tryoni*), the olive fruit fly (*Bactrocera oleae*), and the Mediterranean fruit fly (medfly, *Ceratitis capitata*). The medfly is a highly invasive polyphagous insect and one of the most destructive agricultural pests throughout the world due to its broad host plant range that includes more than 260 different fruits, vegetables, and nuts [14]. The medfly is currently endemic throughout Africa, the Middle East, European countries adjacent and proximal to the Mediterranean Sea, the Hawaiian Islands, the Caribbean, and Central and South America [15]. Thus, the worldwide economic cost due to crop damage, imposed export controls due to quarantine restrictions, and control and prevention of medfly infestation reaches many US$ billions each year. As such, the medfly is the most extensively studied “true” fruit fly at genomic and molecular levels [16] with a published whole genome assembly that is both of good quality and annotated [17].

Control of the medfly is mainly achieved through an area-wide integrative pest management system that includes release of sterilized male flies developed through sterile insect technique (SIT). The current medfly SIT relies on a female-specific genetic sexing strain that carries a yet unknown temperature-sensitive lethal (tsl) mutation which leads to death of female embryos when incubated at elevated temperatures [18]. Resultant males are mass-reared in billions per week for sterilization and released in affected regions. Sterilized males not only control existing medfly populations but also prevent new invasions. Two factors are key to generation of a successful SIT program; i) a genetic sexing strain that enables efficient removal of female flies before sterilization and release of males, ii) a sterilization system that minimizes the fitness cost to the released males. Understanding the dynamics of early embryo development in the medfly could identify new genes to be used in genetic sexing and/or genes that could be targeted for male sterility. Sex chromosomes are key elements in an effective genetic sexing scheme due to their sex-specific presence, and the description of the mechanisms governing their presence is expected to provide a toolkit for advanced SIT applications [19, 20]. Albeit being pivotal over sustainable pest management, insect embryo development remains poorly characterized beyond the model species *Drosophila melanogaster* and the role of long-read RNA-seq in characterizing non-model organisms hasn’t been adequately demonstrated.

Here, we describe the first extensive study of temporal gene expression dynamics during early embryo development in the medfly. We collected single medfly embryos at hourly intervals for the first 15 hours after egg laying (AEL) and performed temporal absolute gene expression quantification of sexed single embryos using long-read RNA-seq. This period encompasses 3 key embryo development stages: the maternal-to-zygotic transition (4-5 hours AEL [21]), cellular blastoderm (∼8.5 hours AEL [13, 22]), and the beginning of gastrulation (9.5-10 hours AEL [13]). We report on the genome-guided transcriptome assembly based on long-read RNA-seq, identification of 3879 new genes, reorganization of maternal transcripts followed by maternal-to-zygotic transition, clustering of genes based on their temporal expression, and zygotic genes. In addition, we provide insights into the presence of dosage compensation on the X chromosome genes of Medfly.

## Methods

### Mediterranean fruit fly rearing

Mediterranean fruit flies (*Ceratitis capitata*) ‘Wiederman’ strain, that were used in this study are a lab strain. Flies were reared in appropriate holding cages at 25 ± 1 °C, 60 ± 10% relative humidity and 14 L:10D cycles according to the conditions previously described [23].

### Embryo collection, RNA extraction and quality control

A schematic of the workflow is shown in **Figure 1A (Supplementary Figure S1)**. To obtain synchronized samples of *Ceratitis capitata*embryos across early development, eggs were collected at hourly intervals from 0 to 15 hours after egg laying (AEL). For each time-point, 20-30 eggs were collected within a narrow 5-minute window to ensure maximal developmental synchrony. Immediately following collection, the 0-hour (time-0) embryos were flash-frozen in liquid nitrogen to halt development. For subsequent time-points, freshly laid eggs were incubated at 25 ± 1 °C for the desired duration and then flash-frozen in liquid nitrogen at the end of each hour. This process was repeated hourly until the 15-hour time-point. Following freezing, samples were transferred to 2 mL tubes containing RNASafeguard reagent (Dominique Dutscher SAS, Bernolsheim, France) to preserve RNA integrity during storage and downstream processing. From each of the 16 time points, we randomly picked 15 embryos which were separately sheared using a pellet pestle cordless motor (Thermo Fisher Scientific) in a 0.5 mL tube. Once homogenized, 2 µL of ERCC[10] Spike-In Mix 1 (Thermo Fischer Scientific) diluted 1740 times in water were added to each sheared embryo and mixed. We initially compared the performance of two kits: Qiagen Allprep spin columns (Qiagen, Germany) and Nucleospin RNA/DNA XS kit (Macherey Nagel, Germany). We chose the Qiagen Allprep spin columns due to higher recovery rate of RNA (90% versus 10%, **Supplementary Figure S2**). From each single embryo we simultaneously extracted total RNA and DNA using the Qiagen Allprep spin columns according to manufacturer instructions. Eluted DNA was used for sexing the embryos. We only analyzed the profile of genomic DNA for embryos at 15 hours AEL. The quantity of the extracted RNA was determined using a Qubit RNA HS Assay Kit (Thermo Fischer Scientific, USA). The quality of the isolated RNA was assessed using an Agilent TapeStation instrument and Agilent RNA ScreenTape kit as per manufacturer’s instruction (Agilent, CA, USA). See RNA metric for each sample in **Supplementary Table S1, Supplementary Figure S3, Supplementary Figure S4, Supplementary Figure S5, Supplementary Figure S6.**

**Figure 1:**
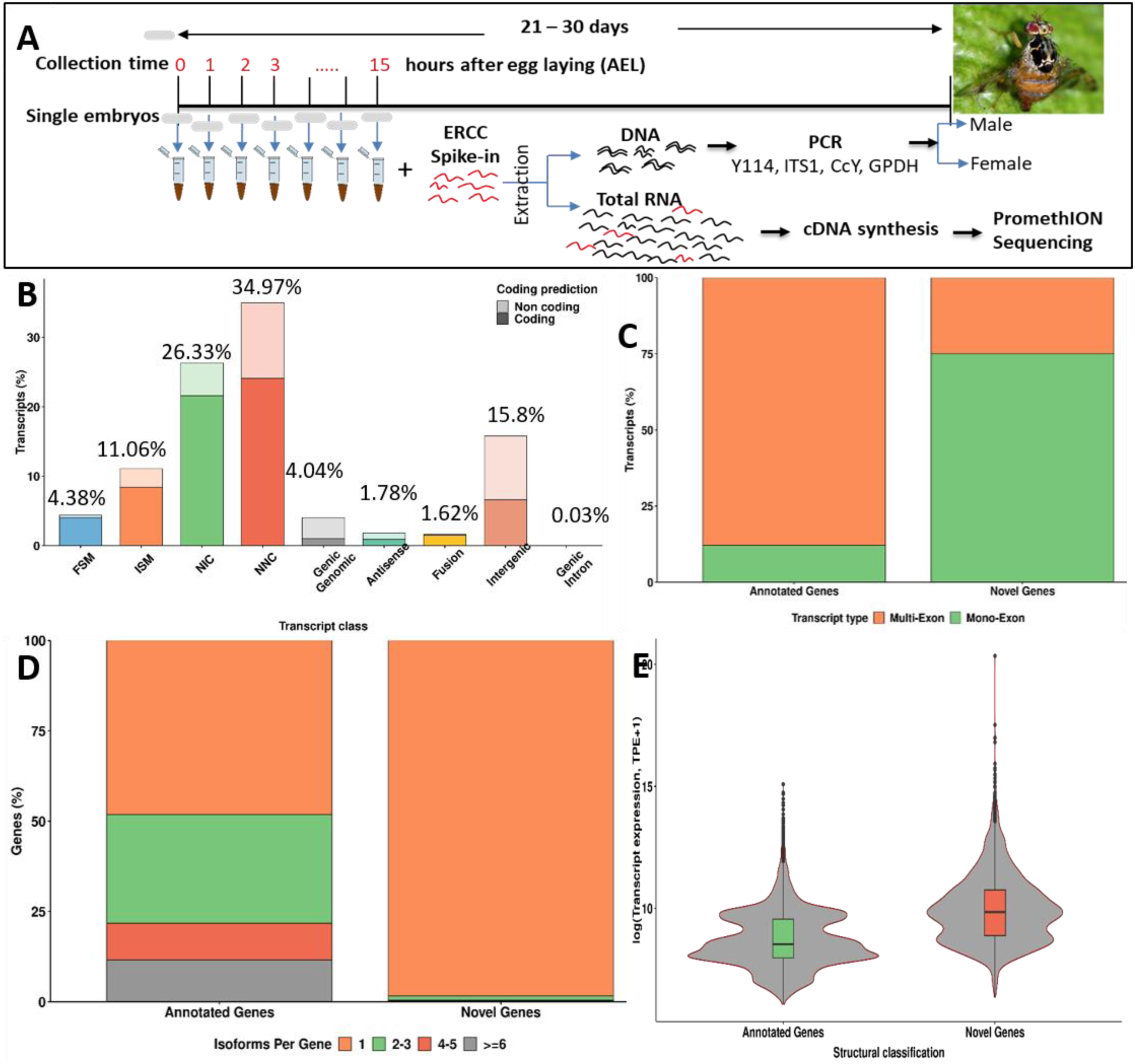
Summary of the workflow and transcriptome assembly. A) Schematic of the library preparation workflow. Single embryos were collected at hourly intervals for the first 15 hours after egg laying (AEL). From each time point, 10 embryos were randomly selected and individually crushed using a pestle and mortar. Equal amounts of ERCC (RNA external standards) were added to each sample followed by DNA and total RNA extraction from each embryo. The DNA was used to sex the embryos via PCR while the RNA was used for cDNA library preparation and long-read RNA-seq. B) Comparison of the long-read derived transcriptome assembly to the NCBI predicted transcriptome. We used SQANTI**[35]** to align, match, and compare gene models in our long-read assembly to the NCBI transcriptome. The results are categorized as. FSM; full splice match, ISM; incomplete splice match, NIC; novel in catalogue, NNC; novel not in catalogue (see **Supplementary Table S1** for explanation). C) Same as B but comparing exon number between annotated and novel genes. D) Same as B but comparing number of isoforms between annotated and novel genes. E) Comparison of gene expression (log transcripts per embryo, TPE) between annotated and novel genes.

### Embryo sexing

We obtained five sets of primers; CcY [11], internal transcribed spacer 1 (ITS1 [24]), Y114 [25], maleness on the Y (MoY, [19]), and glucose-6-phosphate dehydrogenase (G6PD [26]). We optimized the PCR conditions of the primers and also evaluated their suitability to determine the sex of medfly embryos (G6PD was used as a loading control). In our optimized PCR conditions, 12.5 µL of reaction consisting of 1X LongAmp Taq 2X Master Mix (NEB, USA), 0.25 µM of each of the forward and reverse primer, and DNA template were run with initial denaturation at 94 °C for 2 minutes followed by 35 cycles of denaturation at 94 °C for 1 minute, annealing at 58 °C for 1 minute, extension at 72 °C for 2 minutes and a final extension at 72 °C for 5 minutes (**Supplementary Figure S7**). In the end we selected two primer sets for sexing embryos: CcY and Y114. CcY was selected because it yields a different band for males and females. Y114 was selected because females yield no product while males yield a product. These two primer sets also had high sensitivities (**Supplementary Figure S8**). Further, we estimated that enough DNA could be extracted from single embryos to enable sexing by our optimized PCR starting from three hours after egg laying (**Supplementary Figure S9**). We also confirmed a previously reported restriction fragment length polymorphism (RFLP) in medflies **[24]** (**Supplementary Figure S10**). An example of embryo sexing by PCR is shown in **Supplementary Figure S11**.

### cDNA library synthesis

All extracted total RNA was used to separately synthesize cDNA libraries for 15 embryos per time point following our recently published Panhandle protocol [27]. To amplify the cDNA, only 12 of 15 embryos per time point were selected based on their Tapestation profiles and yield. We determined the effect of PCR cycles on cDNA profile (**Supplementary Figure S12**) and used 17 cycles of cDNA amplification for each of the 12 embryos. The quality/profile of each PCR product was assessed using the Agilent 2200 TapeStation system and the Agilent High Sensitivity D5000 ScreenTape. Quantity was estimated using Qubit dsDNA HS Assay kit (Thermo Fischer Scientific, USA).

### Library preparation, sequencing, and basecalling

Ten embryos per time point were selected based on their total RNA profile, cDNA profile, cDNA concentration, and sex, aiming to include equal numbers of male and female embryos. Each sample was barcoded using ONT’s native barcoding kit EXP-NBD196 following manufacturer instructions for the SQK-LSK108 protocol. Samples were then pooled into three batches; time points 0 – 5 hours AEL, 6 – 10 hours AEL, and 11 – 15 hours AEL. A final cleanup of pooled samples was performed using 0.7X AMPure XP beads (Beckman Coulter). **Supplementary Figure S13 s**hows the final Tapestation profile of ready-to-sequence libraries. Sequencing was performed using R9.4 flow cells on the PromethION run by MinKNOW version (3.6.5). All basecalling was performed during the sequencing run using Guppy version 3.2.10. Basecalling included classification of reads as Fail (quality score of less than 9) or Pass (quality score of 9 and above) and assignment to the embryo of origin depending on the barcode sequence added (demultiplexing). Reads whose barcodes could not be reliably identified were grouped as ‘Unclassified’. See **Supplementary Table S2** for read statistics.

### Data processing

A schematic of the data analysis workflow is shown in **Supplementary Figure S14.** Following basecalling, all Pass reads were processed with Porechop to remove sequencing adapters and barcodes followed by Pychopper to classify reads as full-length (containing adapters on both ends), rescued (chimeric reads with barcodes inside the sequence), or unclassified (single or no barcode detected). The resultant reads were combined and processed with Cutadapt [28] to remove their poly(A) tails. Reads were aligned to the NCBI *Ceratitis capitata* annotation release 103 gene models and genome assembly (GenBank GCF_000347755.3, assembly name of Ccap_2.1 [29]) supplemented with ERCC spike sequences. Off-assembly gene sequences were identified by mapping Ccap_2.1-unaligned reads onto the more recent long-read-based assembly EGII-3.2.1 (GenBank accession GCA_905071925.1 [30]). The EGII-3.2.1 assembly is a more contiguous assembly at a scaffold level with scaffold N50 of 77.38 Mb compared to 1.66 Mb for the Ccap_2.1 assembly. However, the Ccap_2.1 assembly is currently the official and annotated release and is more contiguous than EGII-3.2.1 at a contig level with contig N50 of 845 kb compared to 616 kb for EGII-3.2.1.

### Genome guided long-read transcriptome assembly

To obtain a long-read transcriptome assembly of the medfly representative of our samples we aligned adapter and poly(A) trimmed reads from all time points reads to the Ccap_2.1 genome. The resultant aligned reads were processed by FLAIR [31] and Stringtie2 [32]. For FLAIR, samples were processed in 3 batches and their bed files combined before the final ‘collapse’ step.

For Stringtie2, all samples were processed in one batch. Publicly available medfly embryo short-read RNA datasets were obtained from NCBI short read archive and aligned to the genome using STAR [33] to obtain splice junction coordinates. Such junctions with over 3 supporting short reads were supplied to FLAIR to increase splice-site accuracy of the long-reads. The resultant transcripts were further collapsed using ToFU [34] in order to filter out transcripts with less than 90% alignment identity and those with less than 80% alignment coverage and to group transcripts into genes and isoforms based on their alignment coordinates. The resultant transcriptome was analyzed using SQANTI [35]. We filtered the transcripts to remove those with less than 30 long and short reads and those with over 60% adenines downstream of their transcription end site unless they possessed one or more exact splice junction matches to NCBI gene models. ORFinder was used to identify coding sequences (CDS) among transcripts in our assembly. We used custom scripts to either label these transcripts as isoforms of NCBI annotated gene models or label them as independent ‘novel’ genes. Reads that did not align on the Ccap_2.1 assembly were aligned to the EGII-3.2.1 assembly. We repeated the transcriptome assembly process described above. Indeed, some new genes were identified this way and here referred to as off-assembly genes. We generated a new transcriptome assembly and an updated GTF file that included the off-assembly genes, proteome, and transcriptome.

### Pseudo-time analysis

We derived cell clusters using RaceID [36]. The RaceID algorithm was designed for single cell RNA-seq data with molecular counts derived using Unique Molecular Identifiers (UMIs). Since our data lacked UMIs, we instead used absolute gene counts derived using ERCC external standards that were sequenced as part of each sample. Firstly, we removed all genes that were not expressed in at least one embryo. We then filtered out embryos with less than 50,000 total transcript counts. We also removed all genes with less than 1000 transcript in at least one embryo. We then performed lineage tracing using FateID [36].

### Dosage compensation analysis

To assess the chromosome homologies between *C. capitata* and *D. melanogaster*, the gene nucleotide sequences for each assembly were obtained from NCBI. Homologous gene sequences between the two organisms were found with BLASTn [37], filtering for evalue < 5e-4 and percent identity > 60. Syntenic collinear block were then detected with MCScanX (https://github.com/wyp1125/MCScanX) and visualized with SynVisio (https://github.com/kiranbandi/synvisio). To detect dosage compensation on X genes, the EGII_3.2.1 genome was split to its contig component, as this version is known to contain the X and Y sequences in the same scaffold [38]. We then mapped separate male and female short-read datasets available from SRA (ERR4026336, ERR4026337) using bwa-mem [39], selected only unique mapping reads and converted the alignment files to bedGraph using bedtools [40]. We then obtained BigWig files using bedGraphToBigWig (https://github.com/ucscGenomeBrowser/kent). The BigWig files with male and female coverage values were used as input to R-CQ with contig CAJHJT010000034.1.84 as an autosomal normalizer. Contigs having an lnMedian(R-CQ) < -0.4 were selected as X-linked. Embryonic genes were then mapped on the contig sequences using splice-aware minimap2 [41]. We filtered only the primary alignments and selected the genes that mapped on the X-linked contigs. Similarly the genes that mapped on the contigs that comprise scaffold_5 were marked as chromosome 5-linked based on previous chromosomal assignments of the EGII_3.2.1 genome [30]. The normalized embryo counts were assigned to each X and chromosome 5 gene and we excluded those that contained genes of the rRNA cluster that is known to reside both on the X and Y chromosomes of Medfly and genes of the Ceratitis mitochondrion that were found miss-assembled with sequences of the Medfly X. Genes were then filtered based on the consistency of their expression across at least 60% of samples of the same sex that belong to a single developmental cluster. The calculation of expression ratios was performed by dividing the normalized expression values of each X gene with the average expression of chromosome 5 genes.

## Results

### Drosophila melanogaster homolog assignment

The medfly genome is diploid, consisting of six pairs of chromosomes which include a pair of heterochromatic sex chromosomes with the male being the heterogametic sex [42]. The medfly has two draft genome assemblies: Ccap_2.1 (GenBank accession GCF_000347755.3) and EGII-3.2.1 (GenBank accession GCA_905071925.1). The Ccap_2.1 assembly was generated primarily using short-reads[29] while the EGII-3.2.1 was generated primarily using long-reads[30]. Structural and functional genome annotation has been performed for the Ccap_2.1 using the NCBI Eukaryotic Genome annotation pipeline to yield the Annotation release 103. The annotation contains 14,236 genes of which 109 are pseudogenes, 12,563 are protein coding genes while 1,564 are non-protein coding. A total of 4,328 genes have isoforms yielding a total of 22,936 mRNAs. We assigned a *D. melanogaster* homologue to 11,006 (87.6 %) out of the 12,563 protein coding genes (E-value ≤ 1e-3, **Supplementary Table S3**). Of these, 18.7 % were identified in the UniProtKB/Swiss-Prot database, which comprises high quality manually annotated and non-redundant proteins, while the remaining 81.3 % were identified in the UniProtKB/TrEMBL database which contains high quality computationally annotated and classified proteins. Therefore, except for maleness on Y (MoY), all protein names used in reference to medfly are the derived homologs of *D. melanogaster*.

### Medfly *de novo* genome-guided transcriptome assembly identifies new genes and isoforms

We sequenced cDNA libraries from single medfly embryos collected at hourly intervals for the first 15 hours of development including zero time point taken immediately following egg laying. We used our optimized and published Panhandle protocol [27] aimed at capturing poly(A)+ RNA (**Figure 1**A). At each time point we selected 10 embryos for processing. A total of 219 million reads were generated of which 132 million (56 %) were ‘Pass’ and ‘classified’ reads that were used for transcriptome assembly (see **Supplementary Table S2** for sequencing and alignment statistics). Two cDNA libraries separately generated from pooled and mixed-sex embryos at five and six hours after egg laying were also included for transcriptome assembly. We used 3 long-read transcriptome assembly tools; Stringtie2, FLAIR, and Cupcake ToFU to derive our final transcriptome assembly. The *de novo* assembled transcriptome was analyzed using SQANTI [35].

The transcriptome assembly contained a total of 10,740 genes of which 6,861 (63.9%) matched the NCBI annotated genes, while 3,879 genes (36.1%) were new, hereafter referred to as novel genes (**Supplementary Table S4**). The inclusion criteria for novel isoforms and genes is included in Supplementary Materials. The total transcripts identified were 22,857 of which 19,329 were novel, thus, expanding the medfly transcriptome by 85% over the current NCBI annotation. Structural comparison showed that most of the transcripts in the long-read *de novo* transcriptome assembly contained new splice junctions not seen among NCBI annotated genes (NNC, 34.97 %). This was followed by transcripts that contained a new combination of splice junctions already exhibited by NCBI annotated genes (NIC, 26.33 %, **Supplementary Table S4**, **Figure 1**B). Among the 3,879 novel genes that were identified, 1617 (43 %) were predicted to contain an open reading frame suggesting that the rest of the novel genes were noncoding genes. Among the novel protein coding genes, only 14.6 % (236) were assigned a *D. melanogaster* homolog. Further, 75 % of novel genes were mono-exon compared to only 12 % among NCBI annotated genes (**Figure 1**C). Novel genes also contained less number of isoforms (**Figure 1D**) but showed comparable expression compared to annotated genes (**Figure 1E**). This suggested that genes that are non-coding, and/or are mono exon might be more likely to be missed in computational gene prediction pipelines. Noteworthy, the recently discovered mono-exon *Ceratitis capitata maleness on Y* gene (CcMoY[19]), which encodes a key maleness determining protein in medfly was identified among our mono-exon protein-coding novel genes. We further identified 371 genes that were miss-annotated as two or more separate genes but we could find single transcript reads covering these genes (**Supplementary Table S4**, **Supplementary Figure S15**). A more detailed exploration of the long-read derived transcriptome is shown in Supplementary material. We also provide an improved gene model for Doublesex showing that in the medfly, *Dsx* is a 126,100 bp gene composed of 6 exons (**Supplementary Figure S16**). Exon 5 is the key sex-differentiating exon being present in females and spliced out in males leading to a 315 and 394 amino acid protein, respectively, as reported previously [43].

### Unsupervised clustering of embryos traces developmental trajectory

To enable absolute quantification of gene expression, we added equal amounts of ERCC external RNA control mixtures, containing spike-in RNAs of known copy number, to each embryo sample as previously described [9, 44] (**Supplementary Table S5** and **Supplementary Table S6**). This approach, known as absolute quantification or direct absolute normalization, allowed us to estimate transcript copy numbers per gene per embryo. As shown in **Supplementary Figure S17**, this method outperforms conventional relative quantification, especially in transcriptionally dynamic systems like developing embryos. This method was further validated by empirically measured amounts of cDNA generated per embryo (**Supplementary Figure S18**). We further examined pairwise correlation of embryos and show that indeed, adjacent time points had higher Spearman correlation than distant time points, except for the 8 hour time point which showed some anomalies (**Supplementary Figure S19**, **Supplementary Table S7**). Across all our samples, the median limit of detection was 19,000 transcripts per embryo (**Supplementary Table S8**). We then used RaceID and FateID algorithms developed for unsupervised clustering and lineage tracing in single-cell RNA-seq, respectively, to achieve the same objective with our single embryos. A similar application in Drosophila has demonstrated the feasibility and utility of this approach [12]. Although we grouped our embryos into 16 categories based on one-hour sampling intervals following egg laying, RaceID returned 10 clusters (**Figure 2**A). Eleven embryos were excluded from downstream analysis based on the ectopic early expression of previously validated late-expressed genes, such as *sisA* and *slam* [11], as well as other Drosophila-validated markers [45]. These outlier embryos were the sole expressers of these genes within their clusters, suggesting developmental misclassification. Clustering was then repeated, yielding largely consistent cluster assignments and supporting the stability of the grouping (**Figure 2**B). Expression analysis of housekeeping genes confirmed robust transcriptional activity across all embryos (**Supplementary Figure S20**). Embryo sexing showed that as expected, embryos in cluster 1 were mostly not sexed and sexing improved as development progressed (**Supplementary Figure S21**). Lineage tracing showed a single trajectory starting from the least developed to the most advanced embryos (**Figure 2**C). A comparison of real-time and pseudo-temporal grouping showed largely consistent results (**Figure 2**D). For example, cluster 1 contained embryos from 0-3 hours AEL and cluster 9 contained embryos from 11-13 and 15 hours AEL. Nonetheless, certain deviations highlighted the limitations of real-time-based staging. Notably, cluster 10, which was projected as containing the most advanced embryos, contained only embryos collected 14 hours AEL. In this case, the embryos collected 15 hours AEL would have been expected to be the most advanced. These findings highlight the additional resolution gained by transcriptome-based staging over conventional time after-egg-laying classification.

**Figure 2:**
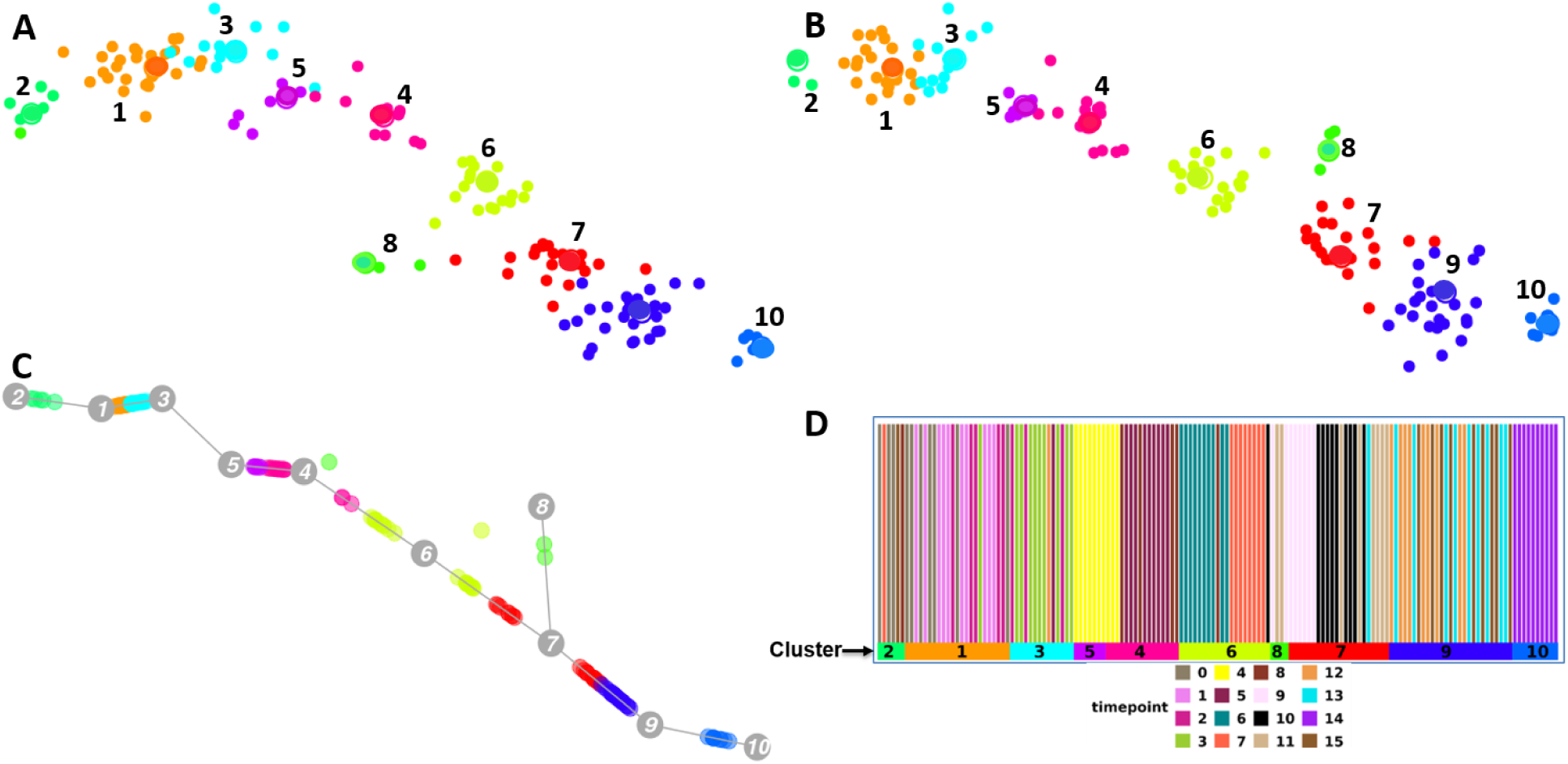
Unsupervised clustering of embryos. A) Unsupervised clustering of embryos using FateID [36]. Clustering was based on normalized absolute number of transcripts per embryo. B) Embryos in ‘A’ were filtered to remove those that had aberrant expression of genes with a previously validated expression profile such as *sisA* and *slam slam* [11], as well as other Drosophila markers [45]. In total 11 embryos were filtered out and the clusters regenerated as in ‘A’. C) Lineage tracing was performed on the clusters in ‘B’ using StemID [36]. The developmental trajectory shows a linear profile from the most immature embryos in cluster 2 to the most mature embryos in cluster 10. D) A schematic comparing pseudo time staging of embryos with staging based on hours after egg-laying (AEL). The embryos are grouped according to their pseudo time clustering with the bottom horizontal bars demarcating the clusters while the vertical bars, which represent each embryo, are colored based on the real-time of each embryo AEL.

### Medfly embryos undergo successive waves of zygotic genome activation

In Drosophila, embryos undergo two waves of zygotic genome activation (ZGA): a minor wave and a major wave. The minor wave, which lasts from about 1 to 2 hours after fertilization [46], is characterized by transcription of about 20 genes by nuclear cycle 9 (NC9) and another 43 by NC10 [47, 48]. The major wave which starts around 2.5 hours after fertilization during NC14 is characterized by transcription of about 3540 genes [47]. We probed the expression of *C. capitata* homologs of these *D. melanogaster* ZGA genes. Out of the 20 early minor ZGA *D. melanogaster* genes, eight had medfly homologs all of which had early expression data (**Supplementary Table S9**). Among these 8 genes, 4 (*sisA*, *CG13712*, *CG6885*, *amos*) were indeed first expressed by embryos in cluster 5 (**Figure 3**A), 3 (*E(spl)m4-BFM*, *l(1)sc*, and *sc*) were first expressed by embryos in cluster 6, while *tsg* was clearly maternally deposited, being expressed by embryos in cluster 1 (**Supplementary Figure S22**).

**Figure 3:**
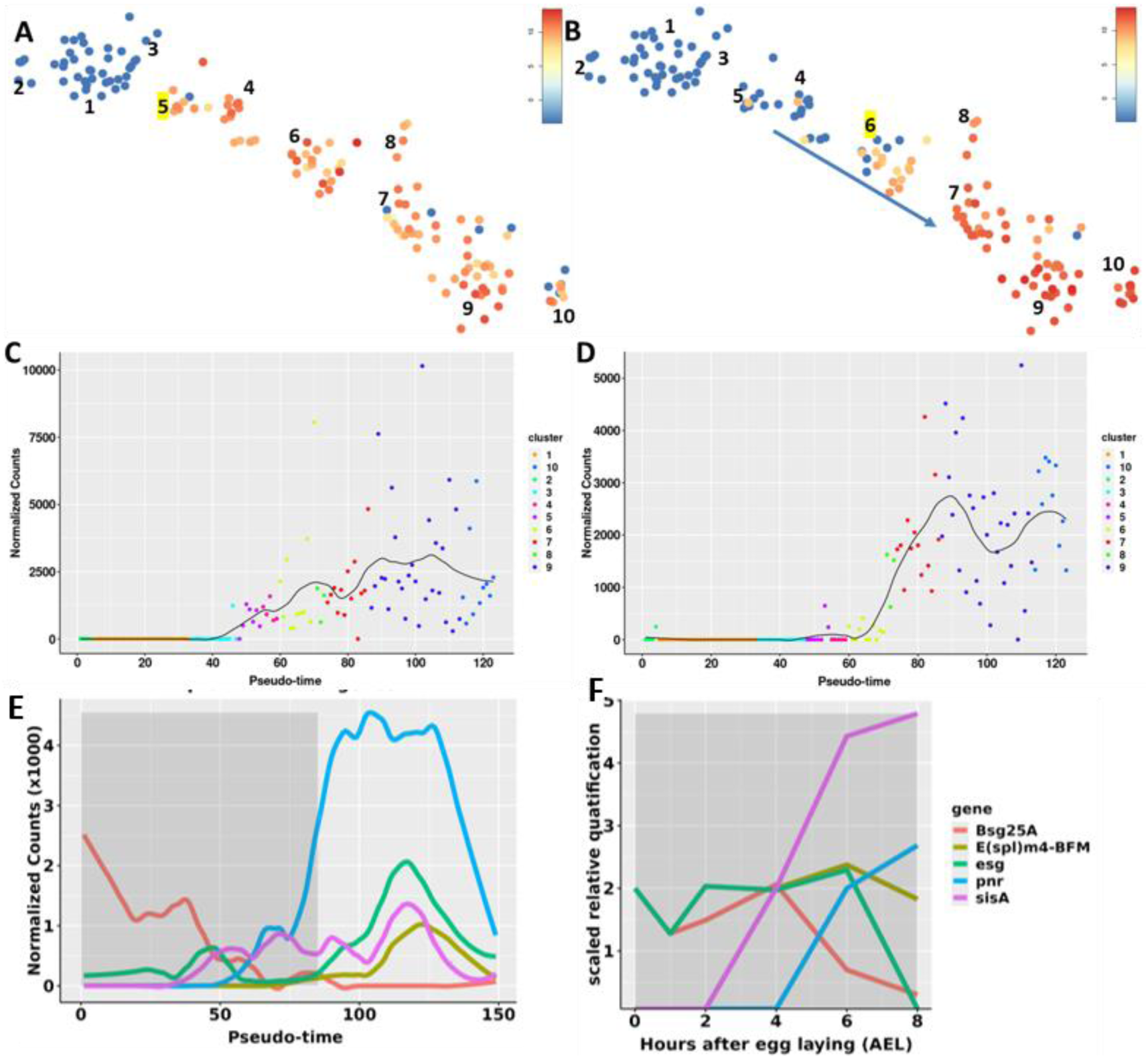
Zygotic genome activation (ZGA). A) We probed the expression of genes previously confirmed to mark the minor wave of ZGA in *Drosophila melanogaster* including *sisA*, *amos*, *CG13712*, and *CG6885* [47]. Cluster 5 showed the first expression of these genes potentially making cluster 5 as a marker of the first wave of ZGA in our medfly embryos. B) Same as ‘A’ except we probed for *D. melanogaster* major wave of ZGA genes including *grh*, *l(1)sc*, *sc*, and *NetA* [47]. Cluster 6 showed the first main expression of these genes potentially marking cluster 6 as a marker of the second wave of ZGA in our medfly embryos. C) Aggregated expression profile of 4 genes used to mark first wave of medfly ZGA (*sisA*, *amos*, *CG13712*, and *CG6885*). D) Aggregated expression profile of 4 genes used to mark the second wave of medfly ZGA (*grh*, *l(1)sc*, *sc*, and *NetA*). E) Expression profile of select genes on a pseudo time scale. F) qPCR validation of the expression of genes in ‘E’. We collected new embryos from 0 to 8 hours after egg-laying and performed qPCR targeting 5 genes (*Bsg25A*, *E(spl)m4-BFM*, *esg*, *pnr*, *sisA*). The grey shaded region shows a similar sampling time between pseudo time and real-time staging.

Among the 43 Drosophila late minor ZGA genes, we had 22 *C. capitata* homologs of which 19 had expression. Among these 19 genes, 9 (47 %) were first expressed by embryos in cluster 5 while 6 (32 %) were first expressed by embryos in cluster 6. Two were expressed in cluster 1 while 2 had sporadic expression in only a few embryos. We therefore, conclude that cluster 5 marks the first wave of zygotic genome activation in our medfly embryos.

Out of the 3542 *D. melanogaster* major ZGA genes, we had *C. capitata* homologs for 2535 genes (72 %). Among these homologs, we had expression for 1477 genes of which 918 were expressed in cluster 1 while 79 were part of the minor wave of ZGA and 147 were indeed, expressed in cluster 6 (**Figure 3**B). We therefore, conclude that cluster 6 marks the second wave of zygotic genome activation in our medfly embryos We then aggregated all genes that were used to mark the two waves of ZGA in medfly and probed their expression on a pseudotime scale. Indeed, genes involved in the first wave of ZGA commenced expression earlier than genes in the second wave of ZGA (pseudotime 40 versus 60, **Figure 3**C and **Figure 3**D, respectively). We further performed qPCR in a new set of embryos collected at different time points to validate the expression of 5 genes known to play a role in early embryo development. qPCR results indeed, validated the profiles we observed using pseudotemporal analysis (**Figure 3**E-F).

We also identified other 334 genes in Cluster 5 that had evidence of being zygotic which could now be new markers of the first wave of ZGA in medfly (**Supplementary Table S9, Supplementary Figure S23**). Cluster 5 was mostly made up of embryos taken at 4 hours AEL. In total, we find 930 genes to be expressed in Cluster 5 and 4 and thus are part of the first ZGA wave in the medfly. Among genes active in the medfly second wave of ZGA, we identified a total of 919 genes expressed in Cluster 6 and 7 that had evidence of being zygotic which we also argue are part of the medfly second wave of zygotic genome activation (**Supplementary Table S9**). These data suggest that medfly embryos undergo two waves of zygotic genome activation clearly marked by clusters 5 and 6, respectively. Clusters 1, 2, and 3 represent embryos that only contain maternally deposited transcripts prior to zygotic genome activation.

Further, we observed that the maternally deposited genes had longer gene lengths compared to zygotic genes (median of 1.8 kb versus 1.5 kb, respectively) although this difference did not reach statistical significance (Wilcoxon p-value, 0.05, **Supplementary Figure S24**). There was no significant difference between the transcript lengths of maternally deposited genes and zygotic genes (Wilcoxon p-value, 0.75). However, the number of exons and total intron lengths of zygotic genes were significantly lower compared to maternally deposited genes (Wilcoxon p-values < 2.2e-16 for both). These data suggested that zygotic genes tend to be short and have less exons.

### Pseudo-temporal clustering highlights relevant biological processes

We performed differential gene expression to determine significantly up or down regulated genes in clusters 1 through 9 (**Figure 4**A). We combined all embryos in clusters 1 and 3 and compared them to embryos in clusters 5 and 4. We then compared embryos in clusters 5 and 4 against cluster 6 (**Figure 4**B). Finally, we compared embryos in cluster 6 against cluster 7 (**Figure 4**C), and cluster 7 against cluster 9 (**Figure 4**D). We then used the significantly down or upregulated genes to perform gene set enrichment analysis using *D. melanogaster* as the reference organism. Relative to cluster 5/4, cluster 1/3 was enriched in metabolic process, mitochondrial gene expression, cellular processes and translation. In contrast, cluster 5/4, where we detected the first expression of Drosophila minor ZGA genes, was enriched in regionalization, pattern specification, multicellular organism development, anatomical structure development, and system development processes.

**Figure 4:**
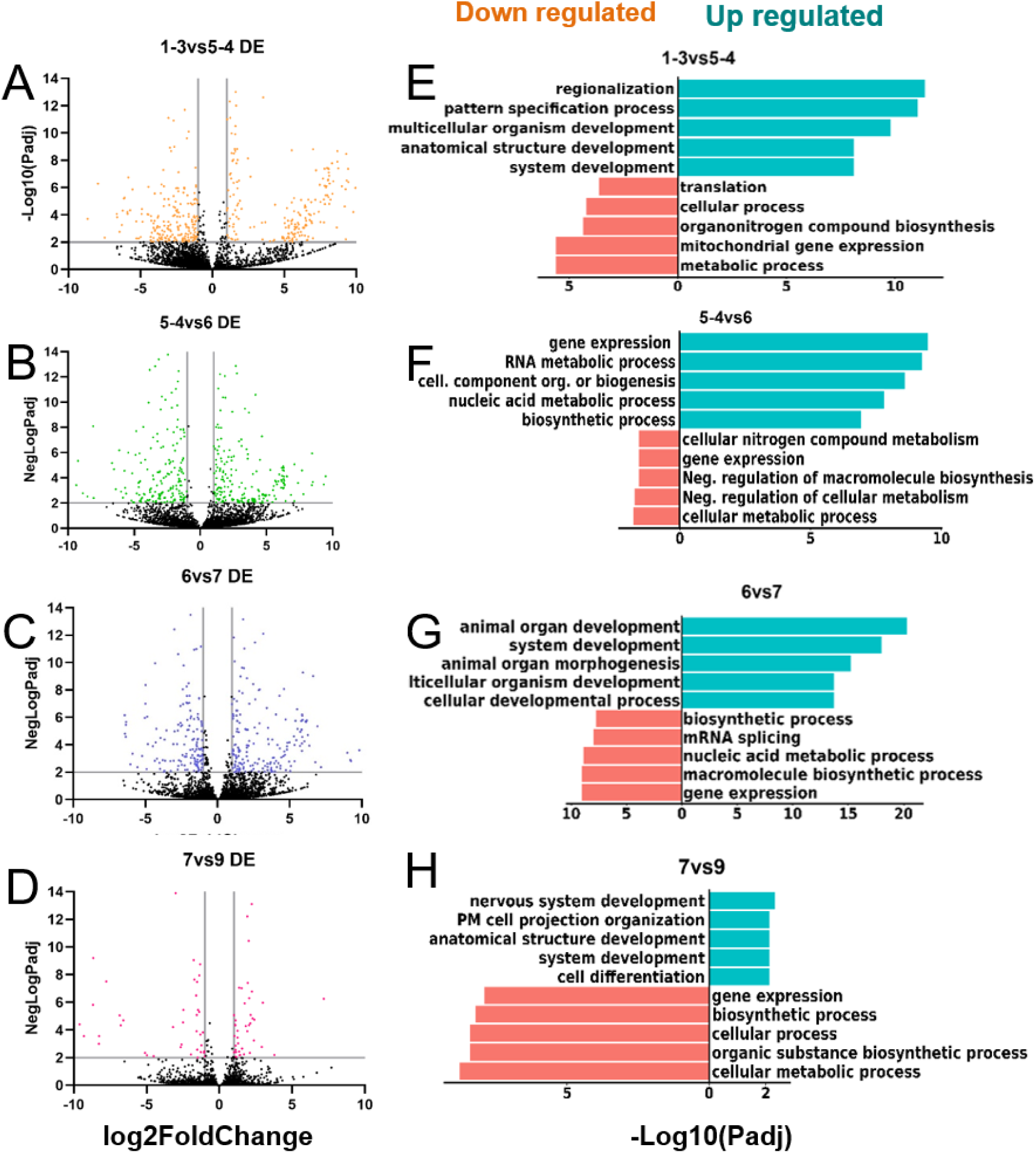
Enriched biological processes in the different clusters. A-D) Differential gene expression was performed in successive clusters. We combined all embryos in clusters 1 and 3. We also combined clusters 5 and 4. We then determined significantly differentially expressed genes in clusters 1-3 versus 5-4 (A), 5-4 versus 6 (B), 6 versus 7 (C), and 7 versus 9 (D). Key genes are shown in each figure. E-H) We used the significantly differentially expressed genes to perform gene set enrichment analysis and sorted the biological process by negative log10 of the adjusted p-value. We picked the top 5 biological processes in each of the clusters. Blue bars to the right indicate significantly upregulated processes while the orange bars to the left are significantly downregulated processes.

Cluster 6, where we first detected the Drosophila major ZGA genes, was enriched in gene expression, RNA metabolic processes, cellular organization, nucleic acid metabolism, and biosynthetic processes. Cluster 7 was enriched in animal organ development, system development, organ morphogenesis, multicellular development, and cellular development process. Cluster 9 was enriched in nervous system development, cell membrane projection organization, anatomical structure development, system development, and cell differentiation Cluster 5 was very interesting as it both showed the first wave of ZGA and was enriched in critical developmental and differentiation processes. We thus probed factors that might be triggering this activation. In Drosophila, the transcription factor Zelda is known to be a major activator of the minor wave of ZGA. The promoter regions of genes transcribed during the Drosophila minor wave of ZGA are enriched in the motif CAGGTAG which is bound by Zelda to mediate the expression of these genes. In the olive fruit fly, we reported a similar finding [44]. However, probing the promoters of genes expressed in the medfly minor wave of ZGA did not return Zelda motifs. However, it is known that many genes do not require Zelda for their regulation during ZGA. Ibarra-Morales et al.,[45] showed that in Drosophila, the transcription start sites of up to 65% of zygotic genes have histone 2A Z (H2A.Z) enrichment. They further showed that knockdown of H2A.Z causes down regulation of a host of genes and compromises 3D chromatic structure. We obtained *D. melanogaster* genes annotated as to whether they contain Zelda motif or are enriched with H2A.Z or not (H2A.Z+ and H2A,Z-, respectively), or have discordant results [45, 49]. For each of the significantly down or upregulated genes in our medfly clusters, we assigned them to their corresponding *D. melanogaster* annotation; Zelda motif, H2A.Z+, H2A.Z-, or discordant. There was an 8 % increase in the Zelda motif enriched genes among cluster 5/4 upregulated genes compared to cluster 1/3 while the number of H2A.Z enriched genes decreased from 80 % to 47 % (**Supplementary Figure S25**). This suggested that neither Zelda nor H2A.Z could explain the mechanism of zygotic genome activation in the medfly. Further into development there was neither clear enrichment of Zelda nor H2A.Z among upregulated genes further suggesting that these mechanisms probably do not play a significant role in medfly zygotic genome activation. The mechanism behind zygotic genome activation in medfly embryos, therefore, remains elusive.

We also used the real-time temporal data and performed temporal gene expressions using DPGP [9, 50] which jointly models data clusters with a Dirichlet process and temporal dependencies with Gaussian processes. We identified 200 gene clusters with differing profiles suggesting specific roles for these clusters during defined developmental periods (**Supplementary Table S10, Supplementary Table S11, Supplementary Table S12**, Supplementary material). We then assigned the clusters to 4 groups (**Supplementary Figure S26**) and found that indeed zygotic genes recapitulate the critical developmental biological processes as seen with the pseudotemporal clustering. Taken together, these results show that cluster 5 marks the switching of key developmental process in our medfly embryos and that the mechanism of this trigger is different in medflies compared to Drosophila with respect to Zelda and H2A.Z.

### Maternal to zygotic transition in medfly shows dramatic reorganization of maternal transcripts

During egg development, the female insect deposits large amounts of proteins and mRNA and other resources into the oocyte. The zygote depends on these maternal transcripts for its early development. The current medfly NCBI gene annotation contains 12,563 protein coding genes. Among these protein coding genes, 2713 (21.6%) were detected at 0 hours AEL (detection limit of 1000 TPE). Among these, the most abundant genes we could assign *D. melanogaster* homologs to were gene7923 and gene6477 which encode Ubiquitin-63E and ribosomal protein L39 and accounted for 1.6 % and 1.1 % of total transcripts per embryo at 0 hours AEL, respectively. We then examined the change in the number of poly(A) transcripts in the developing medfly embryo during its first 15 hours of development. We noticed that the number of transcripts per embryo averaged 0.68 × 10^9^ (±0.14) at 0 hour AEL and then increased to 1.6 × 10^9^ (±0.49) at 1 hour AEL before dropping to 0.46 × 10^9^ (±0.09) at 2 hours AEL (**Figure 5A**). Between 2 hours AEL and 3 hours AEL poly(A) transcripts increased to 1 × 10^9^ (±0.52). We also mathematically converted the number of transcripts per embryo to nanograms of mRNA per embryo and observed a similar trend as would be expected (**Supplementary Figure S27**). By comparison, *D. melanogaster* embryos contain ∼3 × 10^9^ copies of maternal poly(A) transcripts[51] during early development while the closely related *B. oleae* embryos contains 1.5 × 10^9^ poly(A) RNA transcripts at 1 hour AEL[44].

**Figure 5:**
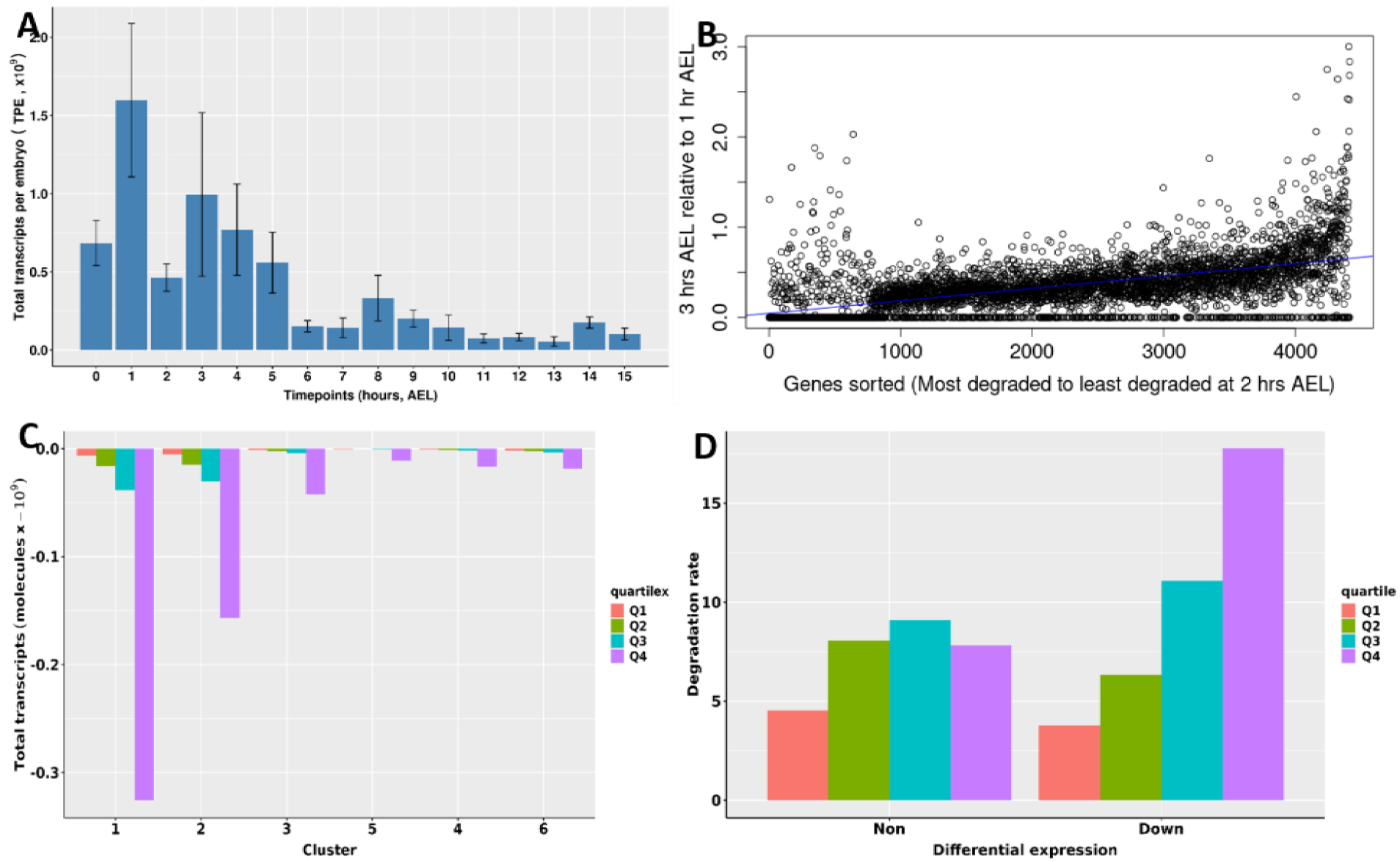
Reorganization of maternal transcripts during maternal-to-zygotic transition (MZT). A) Profile of total transcripts per embryo across time points. We performed absolute normalization to obtain the total number of transcripts per gene per embryo and then summed this. The bars indicate the mean for all embryos at that time point while error bars indicate standard deviation of the mean. B) The ratio of gene expression of embryos at 3 hours after egg laying (AEL) and 1 hour AEL were determined. Genes were sorted based on their gene expression levels at 2 hours AEL, from least abundant (most degraded) to most abundant (least degraded). C) We obtained differentially downregulated genes in cluster 1 relative to cluster 6. These genes were sorted according to their expression levels in cluster 1 and put in 4 quartiles: quartile 1 (Q1) contained the least abundant genes while Q4 contained the most abundant genes. The expression levels of all genes within each quartile were summed across clusters 1 to 6. D) We obtained both differentially downregulated (Down) and non-differentially expressed genes (Non) in cluster 1 relative to cluster 6. These genes were sorted according to their expression levels at in cluster 1 and put in 4 quartiles as explained in ‘C’. We then determine the ratio of expression in cluster 1 relative to cluster 6 to determine the ‘degradation rate’.

We hypothesized that the embryo must be changing the distribution of poly(A) transcripts prior to their maternal-to-zygotic degradation. To test this, we compared the abundance of transcripts at 1 hour AEL relative to the abundance of the same transcripts at 3 hours AEL. We then sorted the transcripts from the most abundant to the least abundant between 1 and 2 hours AEL (**Figure 5B**). We noticed that indeed, there is a re-organization of maternally deposited transcripts such that the least abundant transcripts at 2 hours AEL have a higher abundancy at 3 hours relative to their abundance at 1 hour AEL. We think that this is a deliberate re-organization of maternal transcripts prior to their degradation. Noteworthy, our RNA-seq protocol utilized an oligo(dT) primer for cDNA synthesis and thus, the dynamics observed in the first 3 hours of medfly embryo development relate to poly(A) transcripts. Starting from 3 hours AEL onwards, we observed a gradual reduction in abundance of transcripts per embryo and mRNA from 1 × 10^9^ (±0.52) at 3 hours AEL to 0.1 × 10^9^ (±0.04) at 15 hours AEL, a 90 % drop (**Figure 5A**). We believe that this is a reflection of the degradation of maternal transcripts as part of maternal-to-zygotic transition (MZT). In Drosophila, MZT involves destabilization of up to 35 % of maternally supplied transcripts by maternally encoded proteins by the end of 3 hours after fertilization [52] while in *B. oleae*, we previously detected a 36 % drop in transcripts per embryo by 6 hours AEL.

To elucidate the mechanism of MZT, we returned to the clustering of embryos we performed based on their gene expression. First, we performed a differential gene expression between clusters 1 and 5 and identified significantly down regulated genes. We then ranked the expression pattern of these genes in cluster 1 and put them into four quartiles, Q1, Q2, Q3, Q4, respectively. We then summed the number of transcripts per embryo for all genes in each quartile across all clusters. We noted that the abundance of genes gradually reduced from cluster 1 through cluster 6 (**Figure 5C**). We believe that this is a reflection of the gradual degradation of maternal transcripts as observed in other organisms during maternal to zygotic transition. We then determined the ratio of the total transcripts in each quartile in cluster 1 to that in the respective quartile of Cluster 6, to determine the rate of degradation (**Figure 5D**). Here, we noted a rate (ratio) of 4, 6, 11, and 18 for Q1, Q2, Q3, Q4, respectively. This suggested that the rate of degradation of maternal transcripts was proportional to the initial abundance of transcripts in Cluster 1. Thus, transcripts in quartile 4, which contained the most abundant transcripts in cluster 1 had a 4 fold higher rate of degradation compared to transcripts in quartile 1. On the contrary, transcripts that had shown no significant differential expression showed similar degradation rates regardless of the quartile (**Figure 5D**).

Further, we investigated the presence of X chromosome dosage compensation in our data. We identified scaffolds and contigs in the genome assembly that correspond to X chromosome and autosomes, from which we detected 762 transcripts that derive from the medfly X. We then used the sexing information and pseudotime clustering of embryos to filter X transcripts for consistency of expression across samples of the same sex that belong in the same developmental cluster. This resulted in 380 X genes for which we calculated the per-cluster sex-specific expression and compared it to the expression levels of genes from the autosomal chromosome 5. The results indicate that expression from the X initiates during the minor wave of ZGA at Cluster 5 & 4 and stably increases across the developmental trajectory of the embryos. A comparable ratio of X:Chr5 expression was observed between male and female samples from Cluster 4 and onwards (Figure 6, Supplementary Figure 28), suggesting the presence of dosage compensation on X-linked genes, even from the early onset of transcription activation. No sexing information was available for samples of Cluster 5, thus we cannot exclude a sex biased expression during the initiation of the minor wave of ZGA. Assuming that the chromosome 5 transcript levels represent the diploid expression state, the X gene expression peak at Cluster 9 and 10 indicates that dosage compensation might act towards the direction of the hemizygous state. During Cluster 9, the median X gene expression is almost half that of chromosome 5. During that time-point, females express X genes at ∼52 % of the chromosome 5 levels, with the respective male X expression being at ∼55 %. Similarly for Cluster 10, the female X expression is at ∼72 % with males expressing the X at ∼62 % relative to chromosome 5 levels (**Figure 6**).

**Figure 6:**
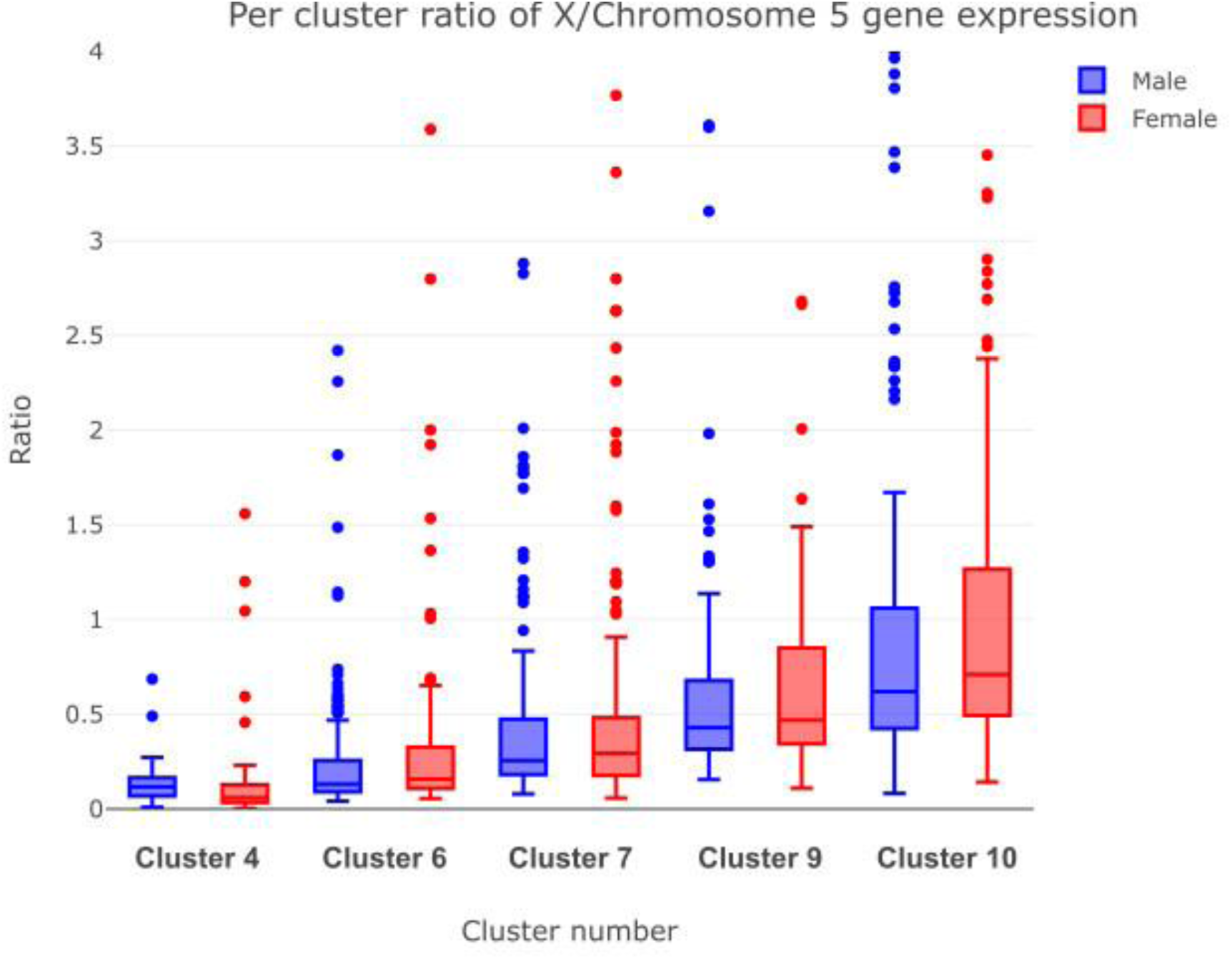
Assessment of dosage compensation in medfly embryos. First, we identified the Medfly early zygotic genes on X chromosome and chromosome 5. We calculated the male and female average zygotic gene expression from chromosome 5 genes across the different clusters. Then, we derived the X:chromosome 5 ratio for each X chromosome gene individually by calculating the average male and female expression of that X gene across different clusters, and dividing it with the respective male or female chromosome 5 average expression.

Taken together, these results suggest that there’s a dramatic reorganization of maternal transcripts in the first 3 hours after egg laying, prior to MZT. Further, MZT happens gradually at a rate commensurate with abundance of transcripts. Finally, the expression of X genes appears to follow the MZT and is compensated early on ZGA between male and female embryos.

## Discussion

The Mediterranean fruit fly (medfly, *Ceratitis capitata*) is one of the most destructive agricultural pests globally, causing significant economic losses due to its broad host range and impact on fruit production. A deeper understanding of its early embryonic development holds great promise not only for advancing basic developmental biology in non-model organisms but also for informing innovative pest control strategies. In particular, early embryogenesis represents a critical window for molecular interventions, as it provides access to gene regulatory networks that could be targeted to disrupt or manipulate development. This knowledge is especially relevant for improving the Sterile Insect Technique (SIT), the most widely used environmentally friendly method for medfly control. SIT relies on the mass rearing, sterilization and release of male flies, which mate with wild females, leading to unviable offspring and a gradual reduction of the population. For SIT programs to be both effective and cost-efficient, it is essential to eliminate females from the production line before sterilization and release. This is achieved using genetic sexing strains (GSS), which incorporate translocations of selectable marker genes such as *white pupae* (*wp*) [30, 53] and *temperature-sensitive lethal* (*tsl*) [54] on the male-limited Y chromosome. The autosomal *tsl* mutation causes homozygous female embryos to die when exposed to elevated temperatures (e.g., 34 °C for 24 hours), enabling selective rearing of males that bear the rescue *tsl* on the Y. Meanwhile, the *wp* mutation, recently characterized at the molecular level [30], renders female pupae white, while the wildtype males pupae remain brown, allowing for automated sex sorting at the pupal stage. Although these mutations have been genetically mapped for decades, only recently have candidate genes for *tsl* begun to emerge (such as *deep orange*, *dor* [55]), providing new molecular insights into this critical trait. Importantly, the earlier females can be removed from the production pipeline, the more efficient the process becomes. Eliminating females during embryogenesis avoids unnecessary expenditure on rearing and expensive larval diets and improves facility throughput. Further, the presence of sex chromosomes is a key element in the sexing scheme with the study of their content and intrinsic mechanisms offering insights and precise alternatives into sexing methodologies. Therefore, characterizing the molecular dynamics of early medfly embryogenesis is not only fundamental from a developmental biology perspective but also strategically valuable for optimizing SIT and identifying novel gene targets for next-generation genetic control technologies.

Here, we used the Oxford Nanopore Technologies long-read technology to study the dynamics of mRNA abundance and transcription within the first 15 hours of medfly embryo development. The study period captures six important developmental events: i) nuclei dispersion along the embryo (0-2.5 hours AEL, stage 1-7 with 1-64 nuclei), ii) syncytial blastoderm (3-3.5 hours AEL, stage 9), iii) appearance of pole cells at posterior end (4 hours AEL, stage11), iv) blastoderm cellularization (8.5 hours AEL), v) appearance of cephalic furrow and start of gastrulation (9.5 hours AEL, stage 14), and vi) initial germ band elongation (10.5 hours AEL) [13].

Our approach targeted poly(A) RNA in single embryos and identified a total of 10,740 genes in the medfly transcriptome. This is comparable to the 9,777 genes reported in 0-3 hour Drosophila embryos [12]. Of these, 63.9% (i.e. 6,861 genes) matched NCBI predicted genes. In our previous work on early embryo development in the olive fruit fly[44], we identified 10,840 genes with a match rate of 83.7%. We identified many more new genes in the medfly compared to the olive fly (3879 versus 1768). Further, we identified a total of 22,857 transcripts in the medfly compared to 78,018 we identified in the olive fruit fly. We implemented more stringent filtering criteria only allowing novel transcripts that had appeared in more than half of the samples. Still, the majority of novel genes identified (75%) in the medfly early embryos were mono-exon. Among these mono-exon novel genes was the recently discovered MoY[19], a gene responsible for determining male sex in medfly embryos. Further, 43% of novel genes were predicted to contain an open reading frame. Again, this list included MoY. We, therefore, contend that although 75% of novel genes we identified were mono-exon, they are true medfly genes and do not represent artifacts. The better annotated *D. melanogaster* genome contains ∼17,000 genes. Therefore, we add 3879 genes to the current 14,236 NCBI genes giving a total of 18,115 genes.

Accurate developmental staging is essential for understanding gene expression dynamics. It is generally accepted that eggs are fertilized during oviposition and, thus, the absolute time following egg-laying has traditionally been used as a standard reference to compare developmental events across experiments. However, some eggs remain unfertilized while others may be fertilized but retained within the female and laid later. These factors can therefore skew timing of biological events. To address this variability, some studies have opted to perform *in vitro* fertilization and use the elapsed time following fertilization as a standard, or manually determine the developmental stage of embryos using morphological features and number of nuclei. Recently, Perez-Mojica et al.[12] demonstrated in *D. melanogaster* that unsupervised clustering of embryos based on transcriptomic profiles, using computational tools developed for single-cell RNA-seq to reconstruct developmental trajectories of cells, provides a quicker, less laborious, and more accurate method to determine the age of embryos. In line with this, we used the unsupervised clustering algorithm, RaceID3[36], which identified 10 distinct clusters. We excluded 11 embryos based on their aberrant expression of *sisA*, *slam*, *tup*, and *NetA,* genes known to be expressed later in embryo development. Gabrieli et al., showed previously that *sisA* and *slam* are not supplied maternally but are encoded by the zygote [11]. Therefore, embryos that exhibited *sisA* and *slam* expression in cluster 1, the earliest cluster, were removed. Lack of detailed molecular characterization in medfly restricted further quality control of the embryos. Nevertheless, removal of these 11 embryos did not drastically alter the clustering.

Using the lineage tracing algorithm, StemID2 [36], we confirmed that the clusters followed a single linear development trajectory, except for cluster 8 which deviated. Ordering the embryos along a pseudotemporal developmental scale showed good overall concordance with realtime ordering. However, embryos from the 8 hour time point showed the most varied distribution with some of these embryos being clustered with the earliest embryos. Overall, we find that unsupervised clustering combined with pseudotemporal trajectory analysis provide an efficient and accurate method to order embryos developmentally, eliminating the need to manually stage embryos or follow strict hourly collections.

Following fertilization, the genome of the zygote remains quiescent until its activation later on in development in a process termed zygotic genome activation (ZGA). In *D. melanogaster*, ZGA occurs during nuclear cycle 14, approximately 1-2 hours after oviposition [46] and is characterized by two waves, a minor wave where about 60 genes are transcribe followed by a major wave involving transcription of at least 3540 genes [47]. We provide strong evidence that medfly embryos also undergo two waves of zygotic genome activation. Gabrieli et al.[21] reached the same conclusion although they found the first wave to start before 4 hours after oviposition while the second wave started 5 after hours after oviposition. In our analysis, embryos clustered differently depending on whether they expressed genes associated with the *D. melanogaster* minor wave of ZGA (cluster 5 and 4) or the major wave of ZGA (cluster 6). A previous study indicated that medfly ZGA starts around 4 hours after egg laying [11]. Our current results support this. Cluster 5, where we detected expression of the first wave of ZGA genes, is mainly composed of embryos collected at 4 hours AEL. In *D. melanogaster*, it has been reported that the minor wave of ZGA coincides with the migration of nuclei to the posterior of the embryo and establishment of pole cells [12, 56]. In the medfly, Stefani et al., [13] reported that at around 4 AEL, pole nuclei arrive at posterior end (stage 11). It is therefore, likely that similarly to *D. melanogaster*, establishment of pole cells coincides with the first wave of ZGA in medflies.

Further, gene-set enrichment analysis showed that cluster 5 was enriched in key developmental processes such as regionalization and pattern specification, providing further indication that these genes are indeed zygotic. The second wave of ZGA in medflies likely occurs ∼6 hours AEL. This is because cluster 6, where we first detected *D. melanogaster* major wave ZGA genes, is mostly composed of embryos collected at 6 hours AEL. We provide a list of genes expressed in the two waves of medfly ZGA. In *D. melanogaster*, the zinc finger transcription factor *Zelda* (*zld*) plays a critical role in timing and activating zygotic gene transcription. The promoters of many early *D. melanogaster* zygotic genes contain TAGteam sites[57] which is principally composed of the motif CAGGTAG. These TAGteam sites are enriched within 500 bp upstream of the TSS of early zygotic genes[57]. Zelda, which is maternally supplied and maintained throughout embryogenesis in *D. melanogaster,* binds the TAGteam sites and acts as a general activator of hundreds of genes during and after MZT, including early sex-related genes such as *sisterless A* (*sisA*), *scute* (*sc*), *sex lethal* (*Sxl*), *deadpan* (*dpn*) and cellular blastoderm formation genes such as ‘slow as molasses’ (*slam*) and *serendipity* (*sry-α*)[58]. Our previous work provided strong evidence that *zld* plays a role in olive fruit fly early zygotic genome activation [44]. In the current work, we did not find the motif CAGGTAG among the medfly early zygotic genes. Gabrieli et al.[21] have also reported lack of TAGteam sites in *Ceratitis capitata* based on *sry-α*. Further, we also found that many well-known *D. melanogaster* early zygotic genes such as *sna*, *kni*, *nullo*, *dpp*, *gt*, *odd*, and *nrt* are maternally supplied in *C. capitata*. Our data suggests that *Zelda* probably does not activate early zygotic genes in *C. capitata* as it does in drosophilids, though it probably does in *B. oleae*[44].

Egg development is arrested at meiosis in many organisms followed by enrichment of the oocyte with maternally supplied transcripts, proteins, and other materials. Past studies in *Drosophila melanogaster*[48] and *Danio relio* (zebrafish)[59], have shown that maternal transcripts represent up to 75 % of total protein-coding genes. Our bulk RNA-seq analysis in the olive fruit fly showed that 62% of protein coding genes were detectable at 1 hour AEL. In the current work, we detected a lower number of protein coding genes of only 21.6% at 0 hours AEL, likely due to insufficient sequencing depth to capture many of the low expressed genes.

We were interested in exploring the dynamics of early embryonic gene expression. In *Drosophila melanogaster* embryos, one of the main development events is the maternal-to-zygotic transition (MZT). MZT involves two main stages: the clearance of a large proportion of maternal transcripts and proteins originally deposited into the oocyte during oogenesis, and the initiation of zygotic transcription[52]. We noted a similar number of transcripts in the early embryo between medfly and olive fruit fly (1.6 × 10^9^ (±0.49) and 1.5 × 10^9^ [44], respectively). However, this is about half the number reported in *D. melanogaster* (∼3 × 10^9^, [51]). It is likely that Drosophila embryos, which develop at about three times the rate of both medfly and olive fly embryos, necessitate larger deposits of maternal transcripts. We also observed a wide difference in the number of transcripts cleared during early embryo development. In Drosophila, MZT involves destabilization of up to 20 % of maternally supplied transcripts by maternally encoded proteins by the end of 2 hours after fertilization (AF) and another 15% destabilized by zygotically encoded protein by 3 hours AF[52]. In *B. oleae*, we previously detected a 36% decrease in transcripts per embryo between 3 and 6 hours AEL. In the current work, we measured a 90% reduction in transcripts per embryo between 3 and 15 hours AEL. It is likely that the shorter timeframe we studied in olive fruit flies and Drosophila explains the difference.

We noticed dramatic changes in mRNA content per embryo that paralleled transcript number fluctuations. The mRNA content per embryo increased 124% between 0 and 1 hour AEL, followed by a 71% decrease between 1 and 2 hours AEL, which rebounded 144% at 3 hours AEL. A very similar pattern was previously observed in olive fruit fly embryo, where mRNA content dropped 51% between 1 and 2 hours AEL and increased 143% at 3 hours AEL compared to 2 hours AEL (the zero hour time point was not included in that study)[44]. Together these findings represent, to our knowledge, the first report of such dramatic mRNA content fluctuations during early embryogenesis in insects. The conservation of this phenomenon in two tephritid species, occurring at the same developmental time post oviposition, suggests that it is a conserved feature warranting further molecular investigation to delineate its mechanism and biological significance. We believe that this dramatic shift in embryo mRNA content precedes MZT initiation and occurs through modulation of the poly(A) tails of transcripts by poly-adenylation and de-adenylation processes. Indeed, in *Drosophila*, the cytoplasmic poly(A) polymerase encoded by *wispy* promotes poly(A)- tail elongation during oocyte maturation and egg activation[60–62], consistent with the observed 124% increase in mRNA content between 0 and 1 hour AEL. Further still, in *Drosophila*, RNA binding proteins that play a key role in MZT, such as *Smaug, BRAT*, and *PUM,* recruit the CCR4-NOT-deadenylase complex to remove poly(A) tails and mediate the degradation of target genes[63–66]. Whether similar or completely different poly-adenylation and de-adenylation events are at play in the medfly remains to be investigated. Following this dramatic event, we noticed a more gradual drop in embryonic mRNA content from 4 hours AEL till the end of our sampling at 15 hours AEL. We believe that this is a reflection of the MZT process initiating at 4 hours AEL. We further explored the manner of maternal transcript degradation by comparing the rate of degradation as a function of initial gene expression level. Our data indicate a gradual degradation process, with the most highly expressed genes having the fastest rate of degradation. This contrasts a report in *D. melanogaster* where the rate of degradation was similar across highly and lowly expressed genes [45]. Our findings suggests that the degradation machinery may act non-specifically, disproportionally affecting the most abundant transcripts, possibly to reduce the levels of the most highly expressed genes as the embryo prepares for cellularization.

Finally, we also provide a profiling of the X-linked Medfly gene expression and the first evidence for the presence of X chromosomes dosage compensation between male and female embryos. Expression from the X is detected early during development, together with the minor wave of ZGA, following an incremental expression pattern across the developmental trajectory. During this, male and female embryos transcribe X-linked genes at the same levels, indicating the presence of X gene dosage compensation. Although dosage compensation has been extensively described in *D. melanogaster*, the same principles do not apply in medfly, or Tephritids in general, as different Muller elements segregate as X chromosome between the two species (Supplementary Figure 7). Our results indicate that dosage compensation in Medfly acts as early during development as the minor wave of ZGA (Figure 6, Supplementary Figure 8), in contrast to the progressive activation in *D. melanogaster* [12]. However, due to limitations in the sexing information we cannot support the compensated expression of the few X genes expressed during cluster 5 (Supplementary Figure 9). In addition, we compared the medfly X gene expression to that of chromosome 5, that is homologues to *D. melanogaster’*s X, for which dosage compensation acts towards the diploid state. We show that in our datasets the expression levels from Medfly’s X reach almost half the level of chromosome 5, suggesting a direction of dosage compensation toward the hemizygous state (Figure 6). Nevertheless, since we observe an increasing X expression trend towards cluster 10, we cannot exclude the further increase of X gene expression up to the dosage compensation at the diploid level.

Our study has some limitations. For example, we did not account for unfertilized eggs and egg-withholding due to lack of enough molecular data to tease these factors out in RNA-seq data. However, we minimized this impact by removing expression-outlier embryos and applying transcriptome-based staging. In Drosophila, egg withholding can vary by up to 10 hours among species [67], highlighting the need for molecular precision in developmental studies.

## Acknowledgements

This study benefited from discussions at International Atomic Energy Agency funded meetings for the Coordinated Research Project “Generic approach for the development of genetic sexing strains for SIT applications”, IAEA (CRP No.: D4.40.03) Vienna, Austria. Particularly, we would like to thank Kostas Bourtzis for discussions and encouragement.

## Author contribution statement

AB, SO, DR participated in study design, conducted the experiments, participated in the data analysis and drafting of the manuscript. DM helped in nucleic acid extraction. AS was responsible for colony rearing and synchronized embryo collections. KDM and JR conceived the study, sourced funding, and oversaw the project execution. All authors read, corrected, and approved the final version of this article.

## Ethics statements

**Studies involving animal subjects:** No ethical approval was required for this study, as the experimental procedures involved invertebrates (*Ceratitis capitata*), which are not covered by animal ethics legislation.

**Studies involving human subjects:** No human studies are presented in this manuscript.

**Inclusion of identifiable human data:** No potentially identifiable human images or data is presented in this study.

## Data availability statement

The datasets presented in this study have been deposited to NCBI’s SRA with project number PRJNA1328737. The individual SRR numbers for each dataset can be found in Supplementary Table 14.

