## Supplementary materials for "Long-read RNA-seq delineates temporal transcriptional dynamics in multiplexed and sexed single medfly embryos"

**Mono exon gene inclusion criteria**

### 1.0 Results

#### 1.1 Sample processing

A schematic of the sample processing workflow is shown in Supplementary Figure S1. Fifteen embryos were selected at each of the 16 time-points and total RNA and DNA extracted. Firstly, we compared the total RNA recovery rate of two RNA extraction kits: Allprep spin column (Qiagen) and Nucleospin RNA/DNA XS kit (Machery-Nagel, Düren, Germany), using samples with known total RNA concentration. We determined that across three amounts of loaded total RNA: 50 ng, 250 ng, and 500 ng, the Allprep spin column had a significantly higher recovery rate with a range of 35 – 90 % compared to Nucleospin RNA/DNA XS kit whose recovery rate ranged 8 - 13% (Supplementary Figure S2). However, Nucleospin kit has a more consistent recovery rate across the tested samples with a coefficient of variation (CoV) of 16.6% compared with Allprep spin column kit whose recovery rate dropped with increasing sample total RNA concentration giving a CoV of 40.5%. We further examined extracted total RNA and noticed that the profiles showed a single peak at ~1.5 kb which contrasts with mammalian total RNA profiles where 2 peaks at ~2 kb and 6 kb are seen representing 18S and 28S ribosomal RNA, respectively. This phenomenon occurs in most insects whose 28S rRNA contains a weak hydrogen bond that easily denatures to release 2 similar sized fragments that run together with the 18S rRNA [1, 2]. Example profiles of extracted single embryo total RNA are shown in Supplementary Figure S3. Supplementary Figure S3 A and B shows profiles exhibited by embryos in very early stages of development. However, we noticed that the total RNA profile of embryos from ten hours AEL onwards had a lower RNA integrity number (RIN) with most samples having profiles which lacked a single peak (Supplementary Figure S3 C and D). We also assayed the profile of DNA extracted from the embryos. Supplementary Figure S4 shows examples of the profiles which not only confirmed extraction of DNA but also suggested that the quality was good enough for PCR. The DNA was to be used to determine the sex of the embryos.

#### 1.2 Validation of PCR conditions for Medfly single embryo sexing

Four pairs of primers (CcMoY, ITS1, Y114, and CcY) that have previously been used to sex medflies were obtained and their performance evaluated. CcMoY [3] targets a 220 bp region within the medfly male-sex determining gene that is located on the Y-chromosome and only found in male insects. ITS1 targets an internal transcribed spacer 1 sequence found in ribosomal DNA [4]. The sequence is found on both X and Y chromosomes; however, presence of a SNP creates a restriction fragment length polymorphism (RFLP) that can distinguish male and female flies. Although the reported PCR products from this primer range from 56 bp – 457 bp the dominant band obtained with our samples was ~800 bp. Y114 [5] targets A-T rich repetitive regions distributed over 90% of the long arm of the Y chromosome [6] and yields a 670 bp PCR product only from male insects [7]. CcY targets a putative long terminal repeat (LTR) retrotransposon MITE insertion on the medfly Y chromosome [8]. PCR amplification results in a single band of 242 bp in female insects while males yield 2 prominent bands; 250 bp and 727 bp. A loading control, G6PD [4] was included to confirm presence of DNA. We obtained the above primers and optimized PCR conditions for these primers using male and female samples of 2 separate medfly strains: Be and EG [9] (Supplementary Figure S7). Our optimized PCR conditions were able to yield the expected bands for each primer set (except for ITS1) allowing us to use these primers in sexing our embryo samples.

#### 1.3 Determining the sensitivity of sexing PCR

We anticipated that the DNA extracted from the earliest time-points would be in very low amounts and thus insufficient for PCR. Therefore, we studied the sensitivity of the different primers that were to be used in the sexing. We performed half-log serial dilution of male DNA starting from 5000 pg/µl to 0.064 pg/µl and used 0.5 µl of the solution as template in a 12.5 µl PCR reaction. CcMoY, which targets a single gene on the Y chromosome, showed the lowest sensitivity; no bands were observed below 8 pg/µl. ITS1 had the highest sensitivity showing bands up to 0.064 pg/µl (Supplementary Figure S8). We then determined the time-point at which we would first expect to have enough DNA to 'sex' the embryos. This calculation relied on the known fact that Tephritidae embryos (*Ceratitis capitata* [10], olive fruit fly [11], Queensland fruit fly [12]) start with a single nucleus that undergoes 4 synchronous rounds of replication per hour (one round every 15 minutes) for the first 4 hours of development, and assuming our DNA recovery rate was 100%. We determined that it would be at 3 hours AEL when the concentration of the extracted DNA would reach 0.8 pg/μL that we would first be able to sex the embryos (Supplementary Figure S9). We also confirmed the presence of the ApoII restriction fragment length polymorphism in our samples by Sanger sequencing on PCR products obtained by using ITS1 primers (Supplementary Figure S10). This can be used to sex embryos that fail with other methods. We decided to use 2 primers for sexing embryos; CcY and Y114. An example of sexing embryos using these two primer sets on 15 embryos from a single time-point is shown in Supplementary Figure S11.

#### 1.4 cDNA synthesis, amplification and RNA sequencing

The total RNA extracted was used in its entirety to synthesize cDNA libraries following our optimized and published protocol termed Panhandle [13]. We determined the profile of cDNA at different cycles as we have previously shown that PCR cycle number can adversely affect these profiles [14]. We found that indeed, 14 PCR cycles and above biased away from long molecules and over-represented short molecules (Supplementary Figure S12). However, in order to obtained ample amplicons for sequencing we amplified our cDNA for 17 cycles. Samples were pooled, sequenced, and basecalled on the PromethION. Supplementary Figure S13 shows the profile of the three pooled sequencing libraries.

#### 1.5 Sequenced read statistics

The data analysis workflow is shown in Supplementary Figure S14. We generated a total of 219.7 million raw reads across all time-points with 75.8% of the reads successfully demultiplexed and assigned to their sample of origin (166 million reads). Further, among assigned reads, 74.4% were classified as PASS while 25.6% were classified as FAIL. Pass reads have a quality value (Phred score) of 9 and above while fail reads have a QC of less than 9. We did not use FAIL reads for any of our analysis. Globally, total raw PASS reads per embryo ranged from 36,000 to 1.6 million per embryo with an average of 0.77 million PASS reads per embryo (Supplementary Figure S28).

There was a generally well-balanced distribution of reads among samples (Supplementary Figure S29). Details of read statistics are included in Supplementary Table 2.

#### 1.6 Alignment statistics

PASS reads from each sample were processed as described above and then aligned to the Cap_2.1 genome supplemented with ERCC sequences and genomic sequences of novel genes that we identified uniquely in the EGII genome and not in the Cap_2.1 genome. Alignment rates ranged from 27 – 72 % with a mean of 48.3%. Alignment to the transcriptome ranged from 45 – 91 % with a mean of 68.6 %. Further, we assessed read alignment to ERCC sequences and mitochondria. Overall, 0.5 – 34% of PASS reads aligned to ERCC sequences with an average of 7.8% while 0.1 – 8.8% of PASS reads aligned to the mitochondria with an average of 1.62%.

#### 1.7 Error correction of reads

Raw Nanopore reads have relatively high error rates compared to other current next generation sequencing technologies. We, therefore, supplied all reads to Canu to perform consensus error correction. Firstly, we compared the read lengths of error corrected reads before and after error correction and determined that no significant changes were introduced in read lengths. Indeed, reads showed very high correlation with Spearman correlation coefficiency of 0.997 (Supplementary Figure S30). Comparison of the error corrected reads before and after error correction showed that the alignment identity was improved from ~70% to > 95% (Supplementary Figure S31).

#### 1.8 Genome guided transcriptome assembly

We aimed to determine the genes and isoforms expressed within the first 15 hours of embryo development. Adapter and poly(A) tail-trimmed reads were aligned to the Cap_2.1 genome assembly. We aimed to have at least 0.5 million reads per embryo aligned onto the genome which was achieved except for 28 embryos out of 150 (Supplementary Figure S32 shows an example from a single time-point). The aligned reads were processed using Flair using the publicly available gene annotation models (obtained from NCBI) for comparison and short-read data to improve splice junction accuracy. We implemented stringent inclusion criteria accepting isoforms with >30 long-read and short-read support, excluding genes with >60% adenines downstream of transcription end sites (TES). We updated the NCBI gene annotation file to include isoforms of genes that were not already annotated and also include the new genes and their isoforms.

#### 1.9 Clustering of genes based on temporal expression dynamics

In rapidly changing systems like developing embryos, precise regulation of temporal dynamics in gene expression is critical for proper development. Therefore, genes with similar temporal expression profiles can be grouped into clusters. Such clusters are thought to have genes with similar biological function [15, 16]. And, since temporal gene expressions have been suggested to follow Gaussian distribution [17], we used DPGP [18] statistical tool which jointly models data clusters with a Dirichlet process and temporal dependencies with Gaussian processes. We identified 200 gene clusters with differing profiles suggesting specific roles for these clusters during defined developmental periods (Supplementary Table 10). We then assigned the clusters to 4 groups namely: i) Maternal-upregulated: comprising genes whose abundance increased between 0 - 1 hour AEL and were downregulated at 2 hours AEL and generally decreased after 3 hours AEL. ii) Maternal-downregulated: genes whose expression was downregulated from 0 hour AEL and generally stayed low. iii) Transient: genes whose expression peaked only at specific time points, and iv) Zygotic: genes whose expression was only detectable starting from 4 hours AEL implying that they emanated from zygotic genome. A representative cluster is shown for each of the groups in Supplementary Figure S26. A heatmap is also shown combining all genes in each group (**Error! Reference source not found.**B). We performed gene ontology (GO) enrichment analysis (Supplementary Figure S26C) to identify enriched biological process in each of these four groups. Maternal-upregulated genes, were enriched in cellular metabolic processes, nitrogen compound metabolism, gene expression, translation, peptide and amide biosynthesis processes among other processes (Supplementary Figure S26C). Maternal-downregulated genes were enriched in melanin and secondary metabolic processes, regulation of immune system, regulation of catalytic activity, regulation of protein activation, among other regulatory processes. Transient genes showed enrichment of anatomical structure development, developmental processes, cellular process, system development, cell development, cell differentiation, response to stress, among other processes. Strikingly, Zygotic genes were enriched in specific and key tissue formation and developing processes including animal organ development, system development, neurogenesis, cell differentiation, tissue development, sensory organ development, pattern specification process, cell fate commitment, among other similar processes. Supplementary Table 11 lists all clusters in a particular category while Supplementary Table 12 provides all gene ontology enriched processes in each cluster.

### Supplementary Figures

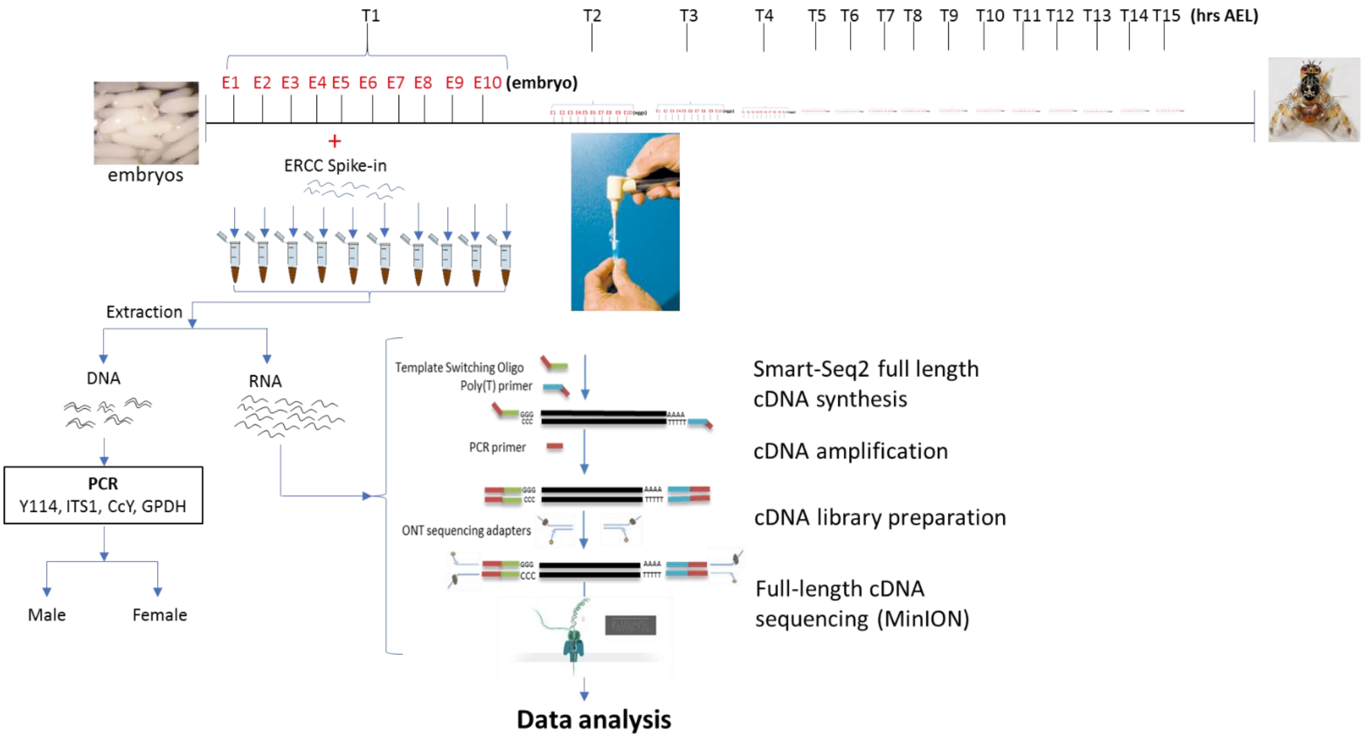


**Supplementary Figure S1: Schematic of the library preparation workflow.** Single embryos were collected at hourly intervals for the first 15 hours after egg laying (AEL). From each time-point, 10 embryos are randomly selected and individually crushed using a pestle and mortar. Equal amounts of ERCC (RNA internal standards) are added to each sample. DNA and total RNA are separately extracted from each embryo. The DNA is used to sex the embryos while the RNA is stored for later use in library preparation and long-read RNA-seq.

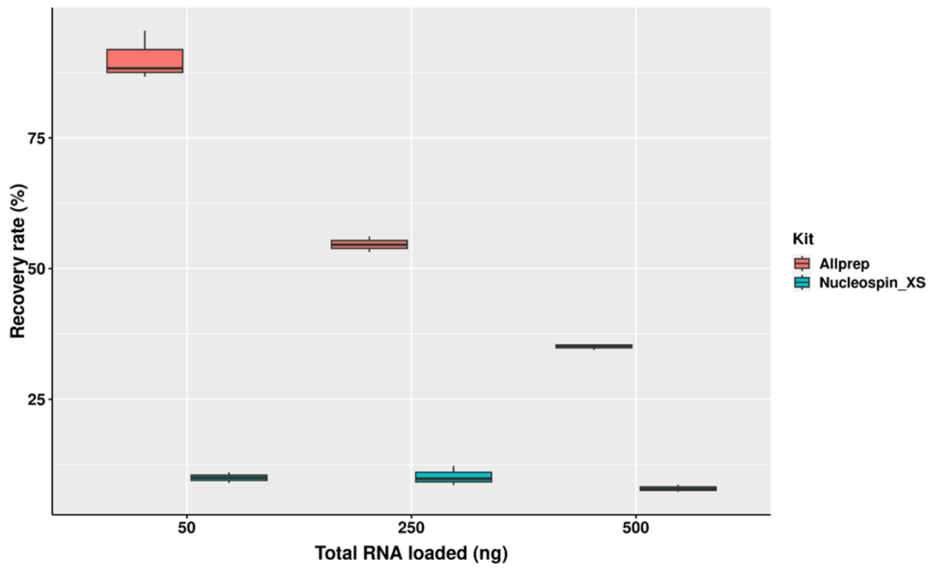


**Supplementary Figure S2: Three technical triplicates of known amounts of total RNA were taken through the total RNA extraction and purification protocol.** The amount of total RNA in the original and first eluted fractions were determined using Qubit High sensitivity RNA kit.

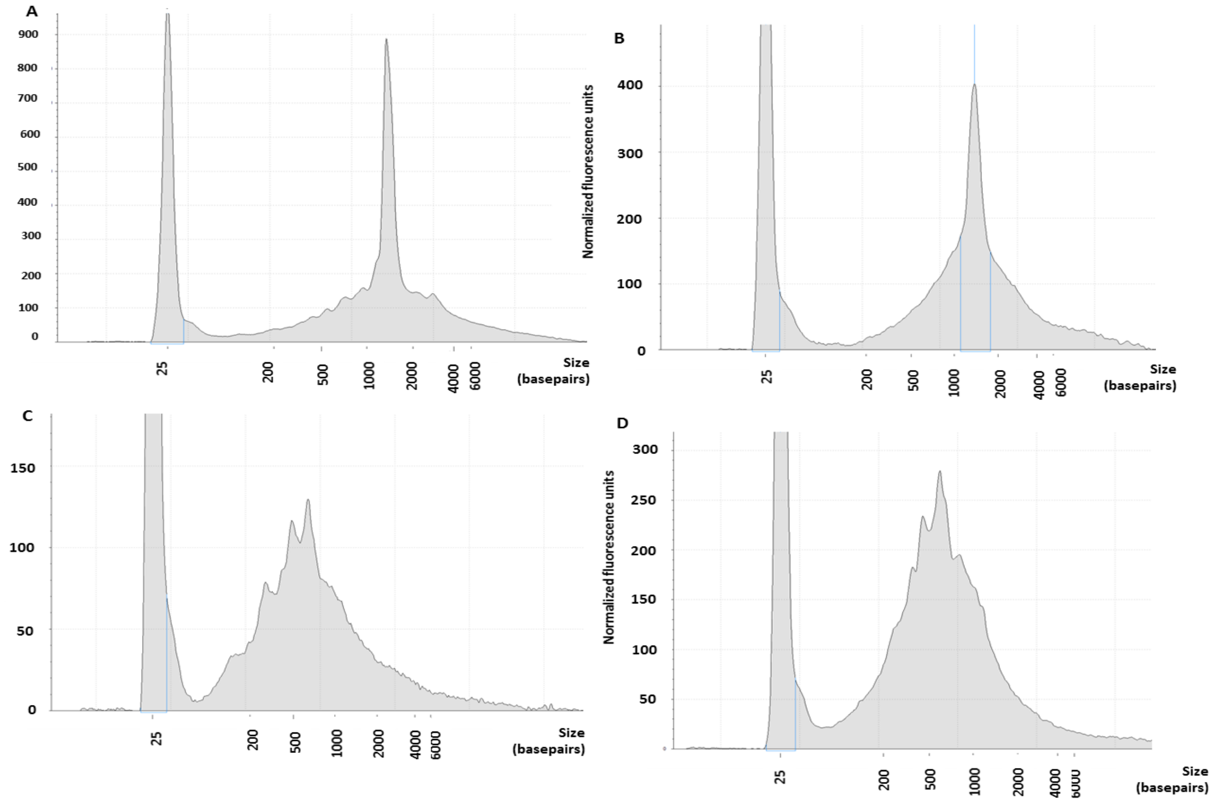


**Supplementary Figure S3: Electrophoresis profile of total RNA extracted from Medfly single embryos**. Single embryos were homogenized using a pestle and mortar. ERCC internal spike-in were added to each sample and the RNA extracted using NucleoSpin RNA XS column. The RNA quality and profile were assessed using eukaryotic RNA ScreenTape. Panels A-D represent 4 different embryos, respectively.

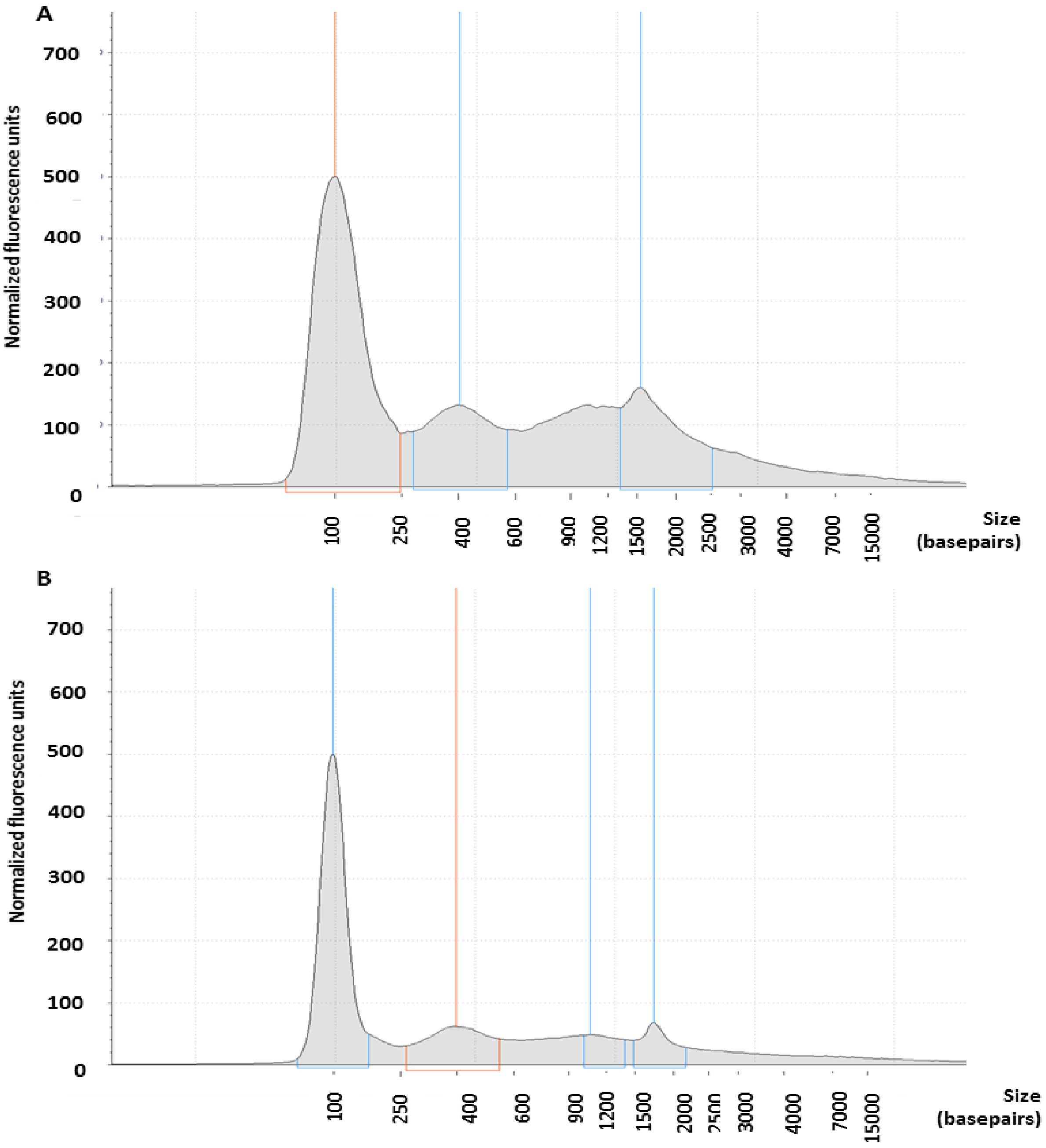


**Supplementary Figure S4: Tapesation profile of genomic DNA.** Genomic DNA was extracted following the Nucleospin protocol (Machery Nagel). The profile of eluted DNA was assessed using Tapestation. A and B show single embryos from the 15 hours after egg laying time-point.

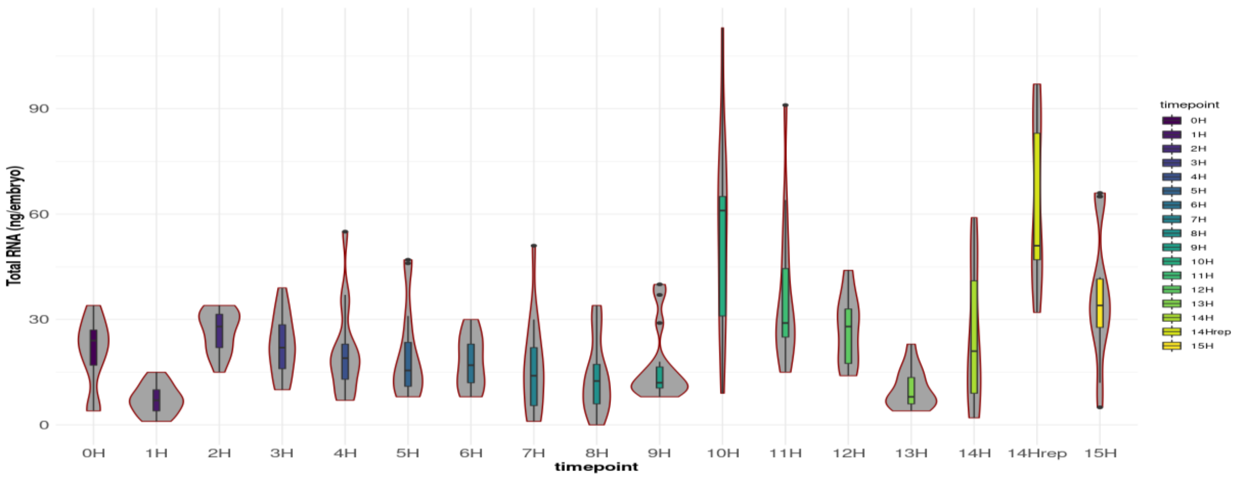


**Supplementary Figure S5:** Total RNA yields across time-points. Total RNA was extracted using Allprep spin column (Qiagen) and the quantities determined using Qubit RNA HS Assay Kit (Thermo Fischer Scientific).

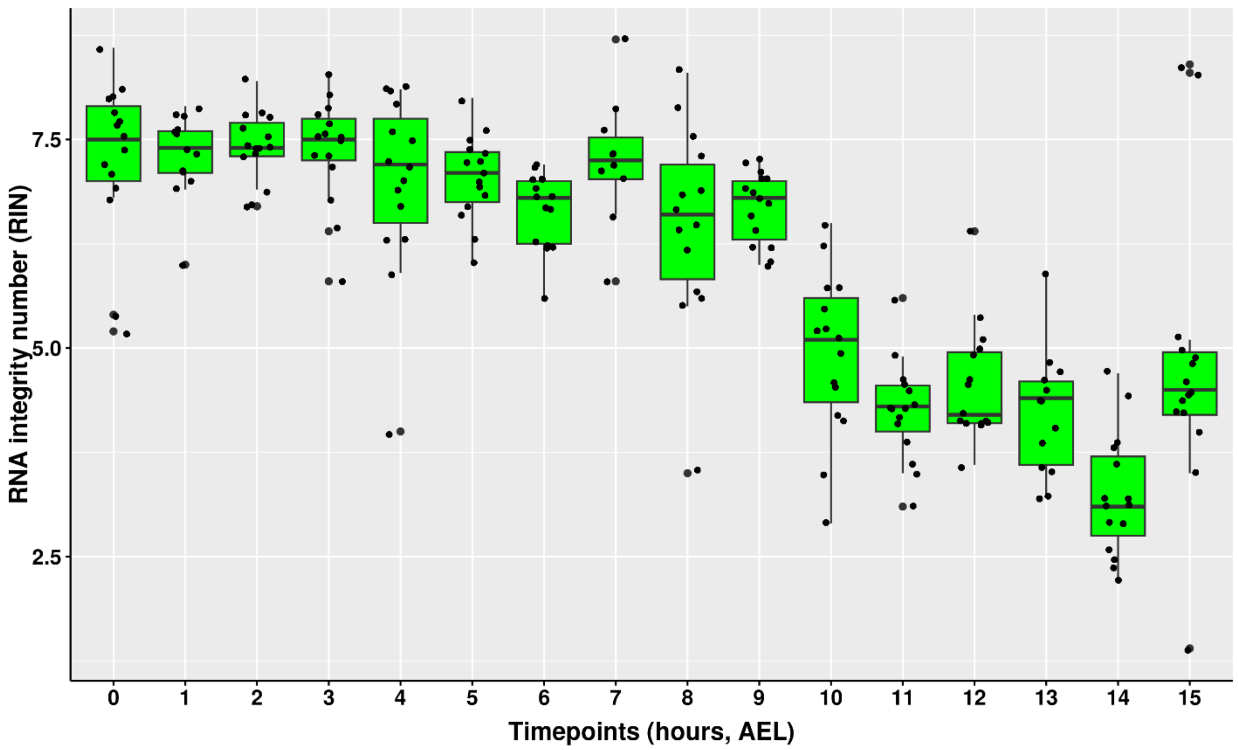


**Supplementary Figure S6:** Same as Supplementary Figure S5 but showing the RNA integrity number (RIN).

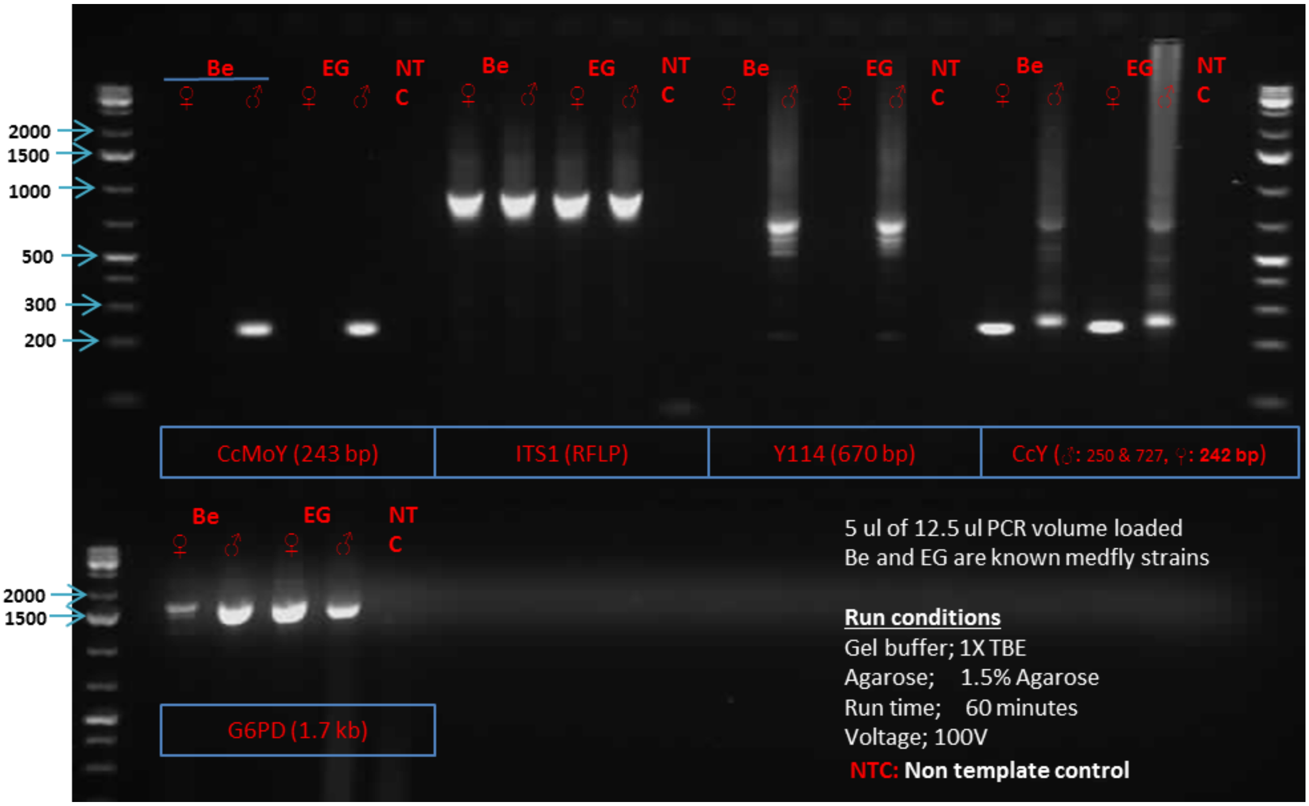


**Supplementary Figure S7: Validation of conditions for Medfly PCR molecular sexing.** Male and female genomic DNA from 2 strains; Be and EG was used as template in PCR. Five primer sets were tested; CcMoY, ITS1, Y114, CcY, and GAPDH. The resultant PCR products were run on a 1.5% Agarose gel (1X TBE, 100V run for 1 hour). The leftmost and rightmost lanes show 1kb ladder.

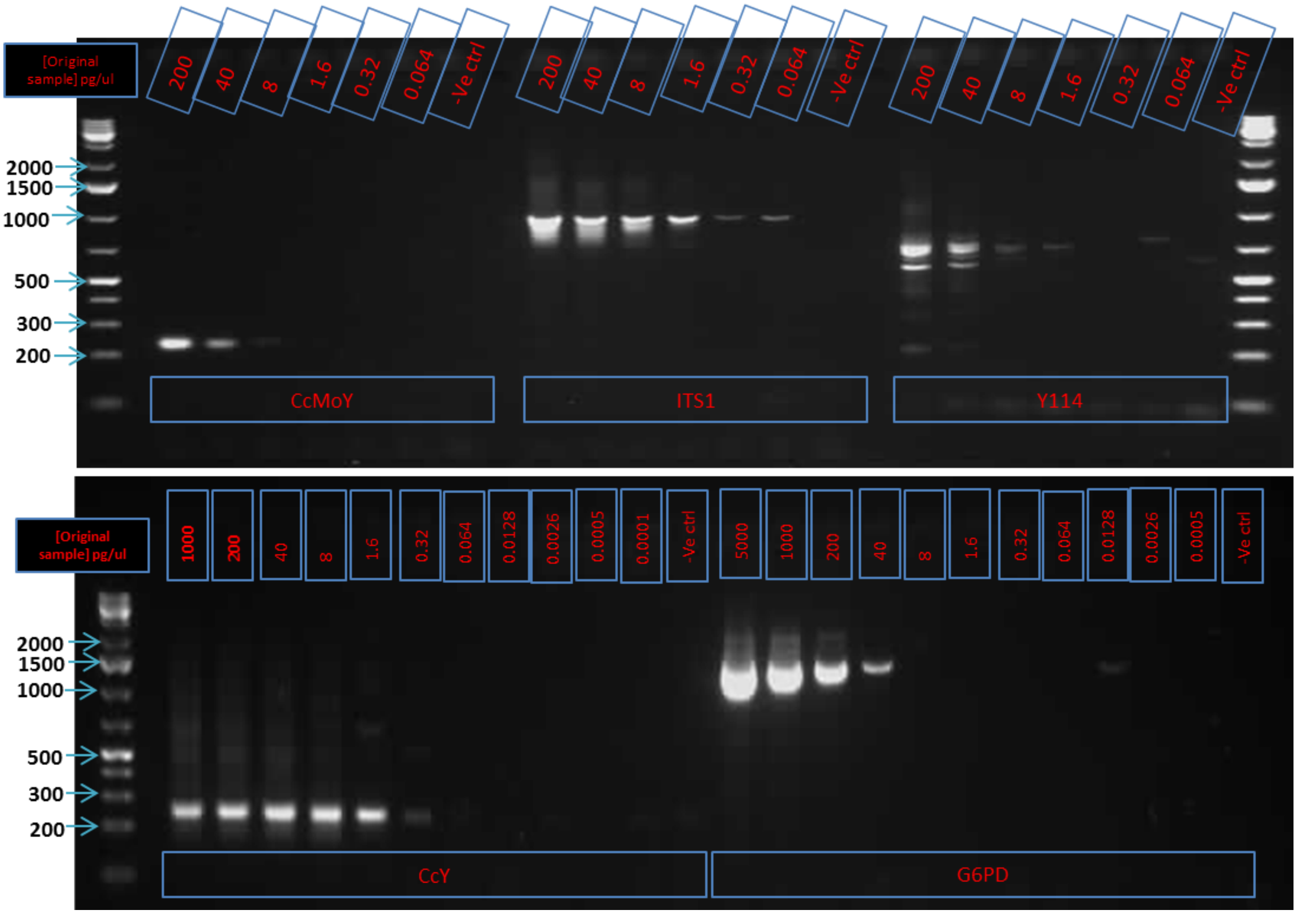


**Supplementary Figure S8: Determining the sensitivity of the sexing-PCR.** Male Medfly DNA was serial diluted starting from 5000pg/µl to 0.064 pg/µl and 0.5 µl used as template in a 12.5 µl PCR reaction. Five primer sets were tested; CcMoY, ITS1, Y114, CcY, and GAPDH. The resultant PCR products were run on a 1.5% Agarose gel (1X TBE, 100V run for 1 hour). The leftmost and rightmost lanes show 1 kb ladder.

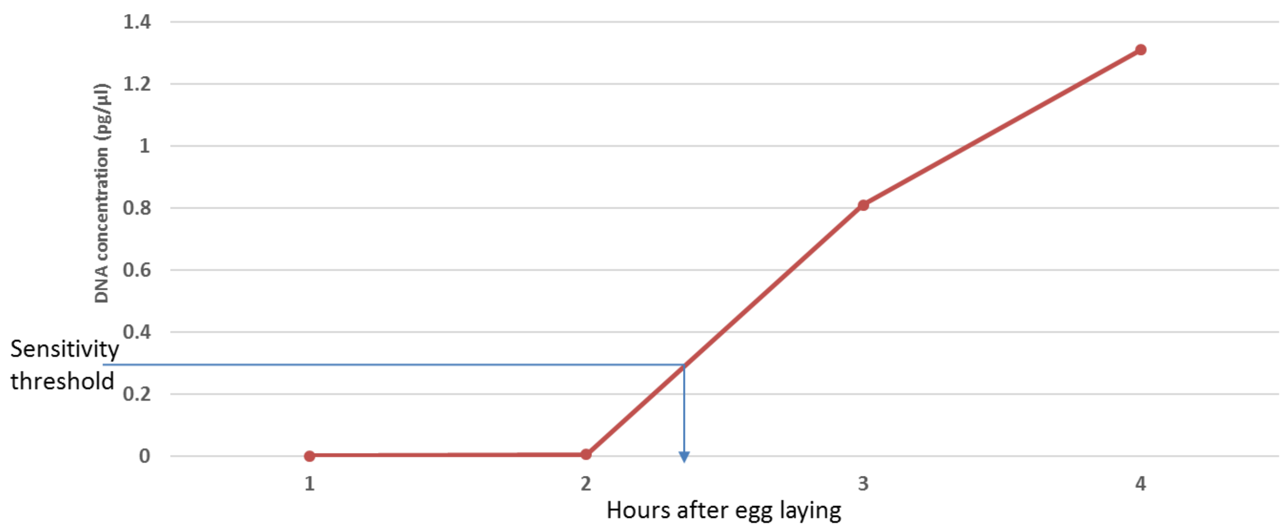


**Supplementary Figure S9: Estimation of expected DNA concentration from single embryos at different time-points.** Since Medfly embryos undergo synchronous rounds of DNA replication within the first 4 hours of development and given the 80 µl elution volume of the DNA extraction kit we used, we estimated the concentration of the extracted DNA assuming 100% recovery at the different time-points. A sensitivity threshold based on CcY primer in Supplementary Figure **S6** above was set at ~0.32 pg/µl.

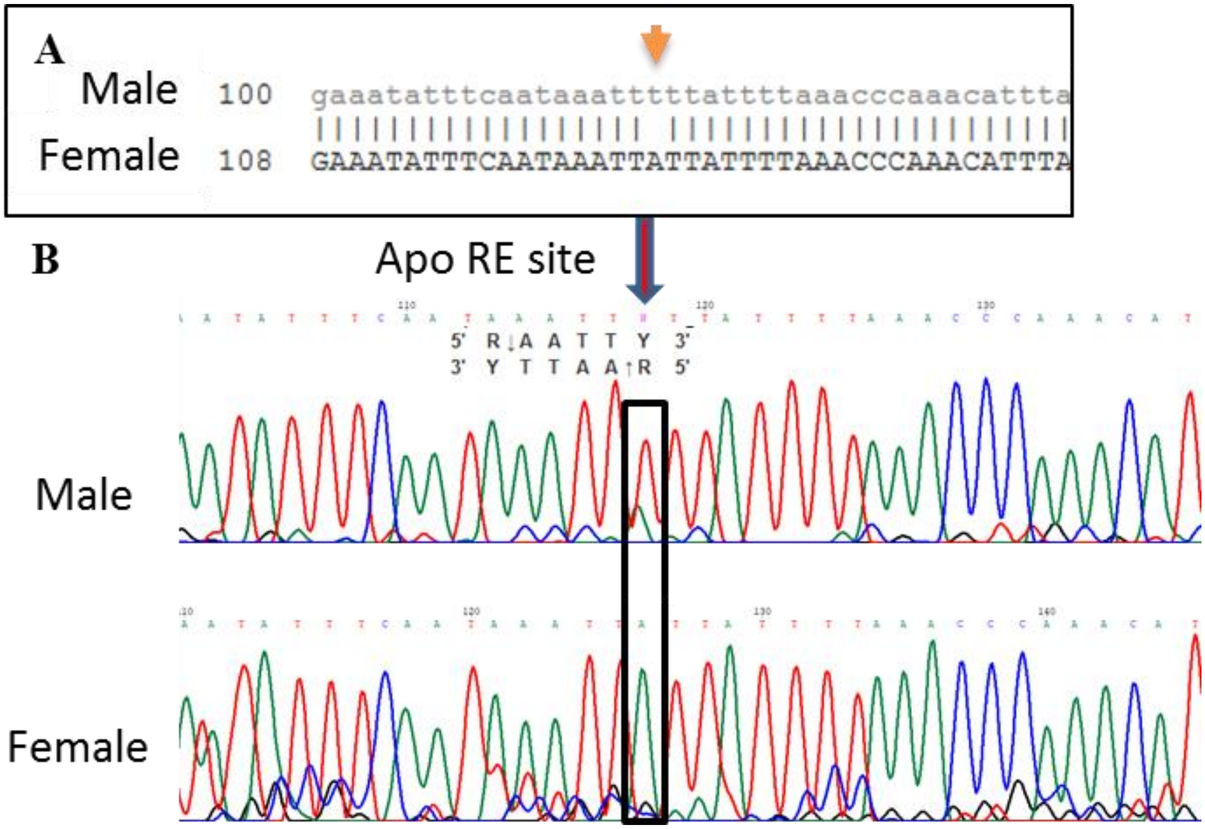


**Supplementary Figure S10: Confirmation of a restriction fragment length polymorphism (RFLP) in our flies.** A) Sanger sequences of a male and a female sample Medfly embryo which confirmed presence of a RFLP as previously reported [4]. B) Electropherogram of the sequences showing the heterozygosity in males and homozygosity in females as previously reported [4].

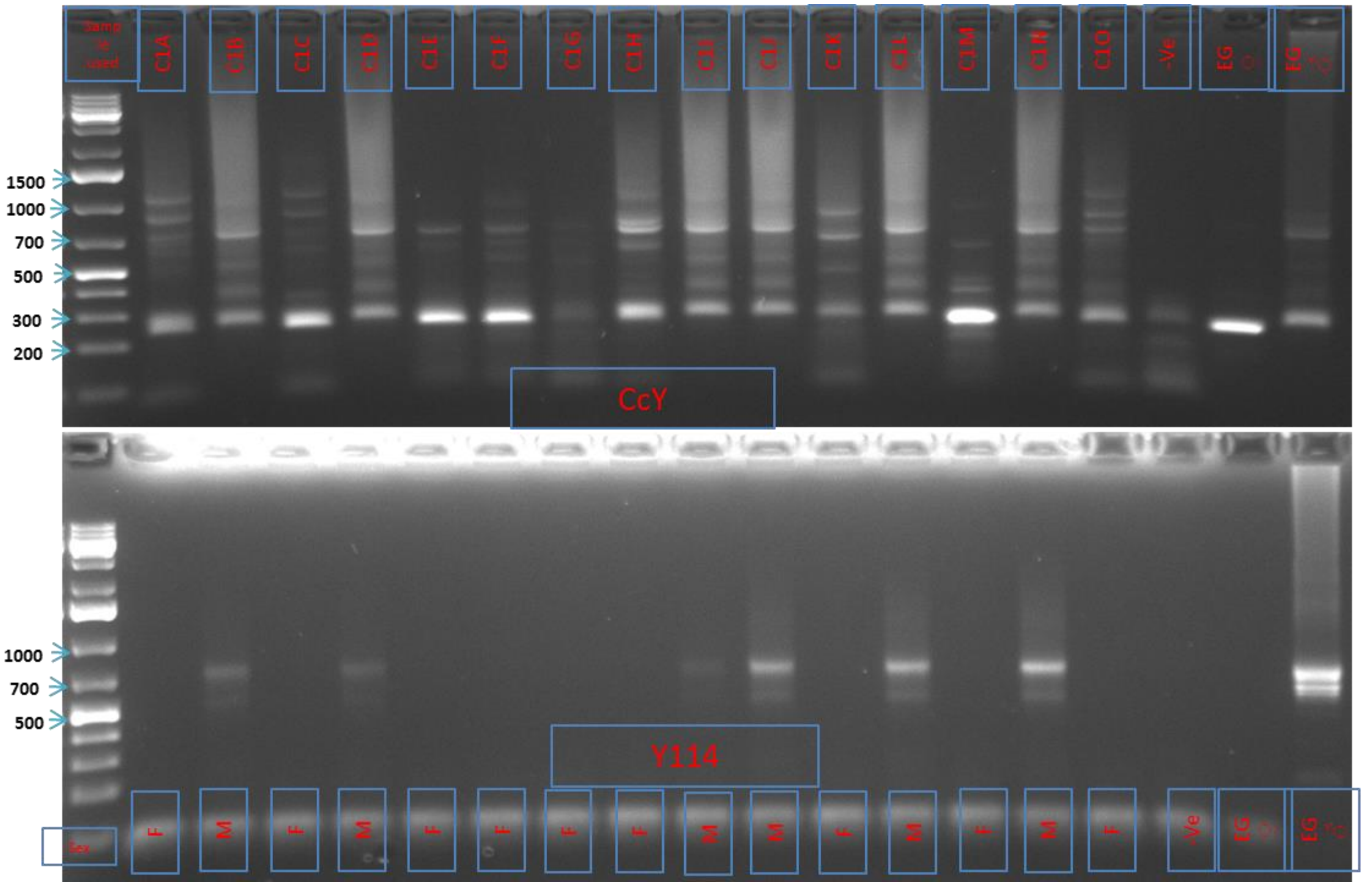


**Supplementary Figure S11: Molecular sexing of Medfly single embryos.** Genomic DNA was extracted from single embryos using NucleoSpin RNA XS columns. The DNA was used as template in 2 separate PCR reactions using 2 sets of sexing primers; CcY and Y114. The resultant PCR products were run on a 1.5% Agarose gel (1X TBE, 100V run for 1 hour). The leftmost lanes show 1kb ladder while the rightmost 3 lanes show non-template control (NTC), Medfly female DNA and Medfly male DNA, respectively. The inferred sex of the embryo is represented as F (female) or M (male) on the bottom gel having the Y114 primers.

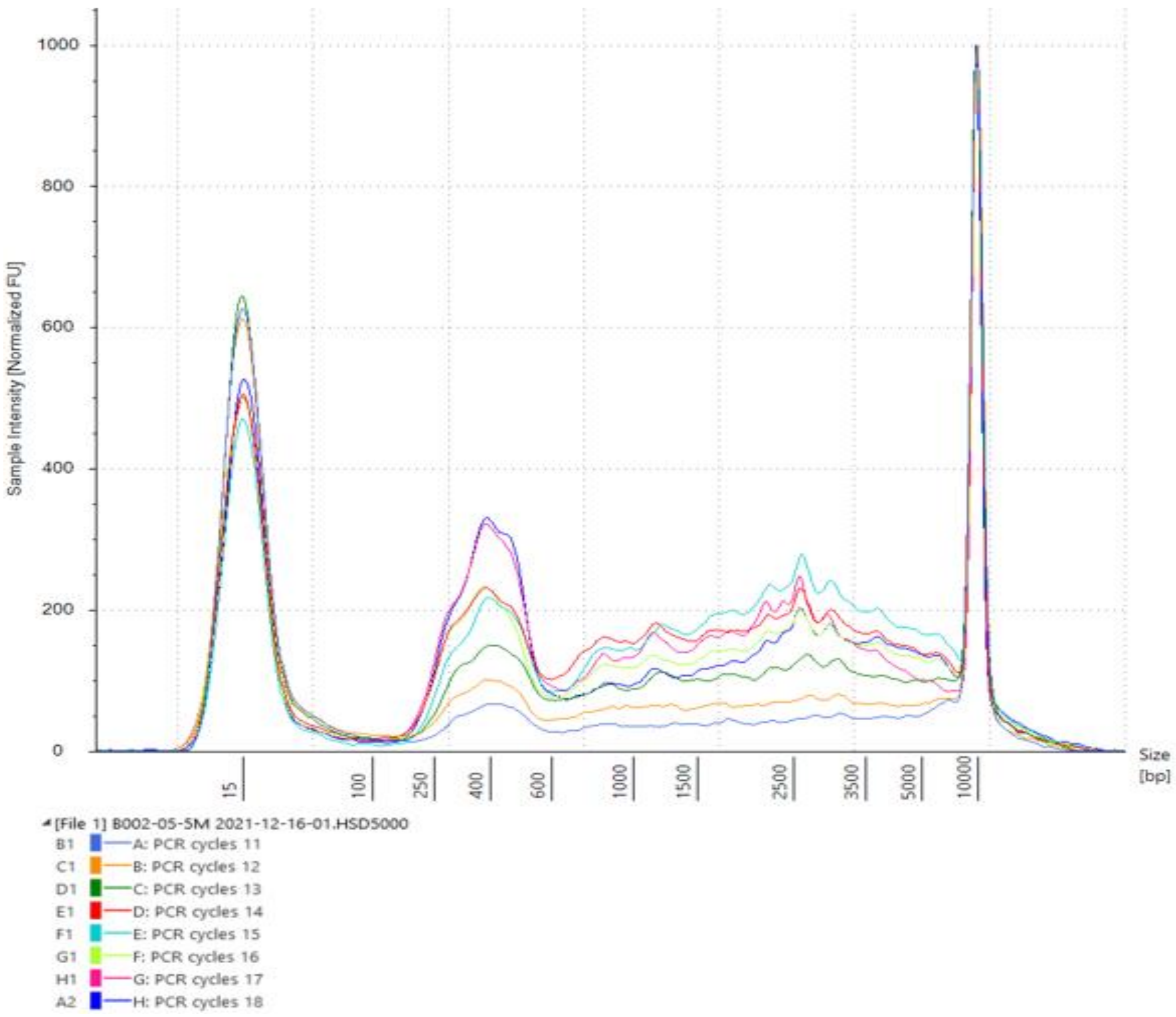


**Supplementary Figure S12: Determination of optimal PCR cycle number.** Tapestation profile of amplified cDNA showing the profile of cDNA following different rounds of PCR amplification. A) shows 15, 16, and 17 rounds of amplification while B shows 8, 10, 12, 13, and 14 rounds of amplification.

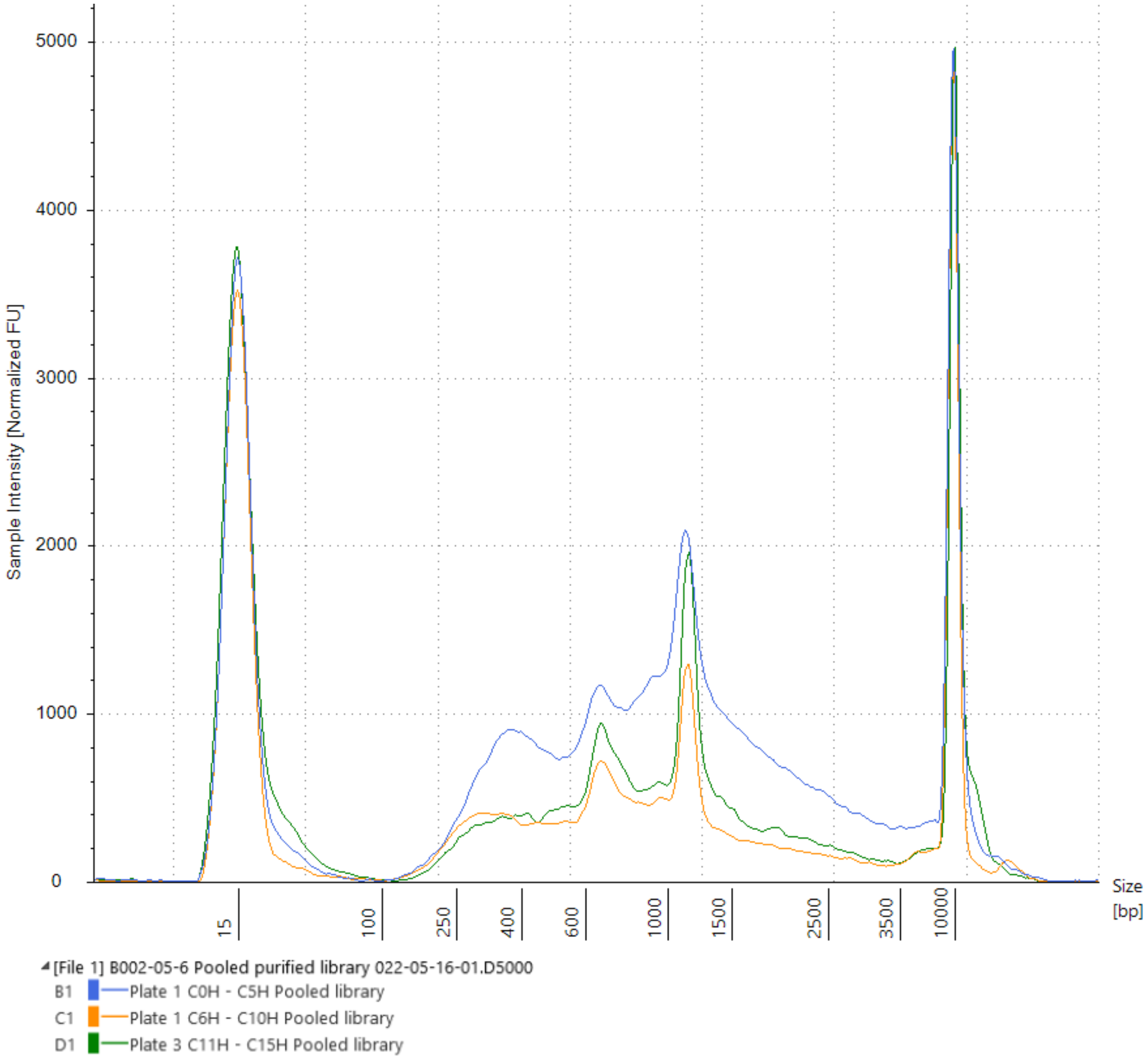


**Supplementary Figure S13: Profile of the three cDNA sequencing libraries.** We selected 10 samples for each of the 16 time-points for cDNA library preparation following our published protocol[13]. Each of the samples was then barcoded using ONT's native barcoding kit, EXP-NBD196, following manufacturer instructions. Samples were then pooled in three batches, time points 0 – 5 hours AEL, 6 – 10 hours AEL, and 11 – 15 hours AEL. The profile of the three purified cDNA sequencing libraries was determined using Agilent High Sensitivity D5000 ScreenTape and is shown here.

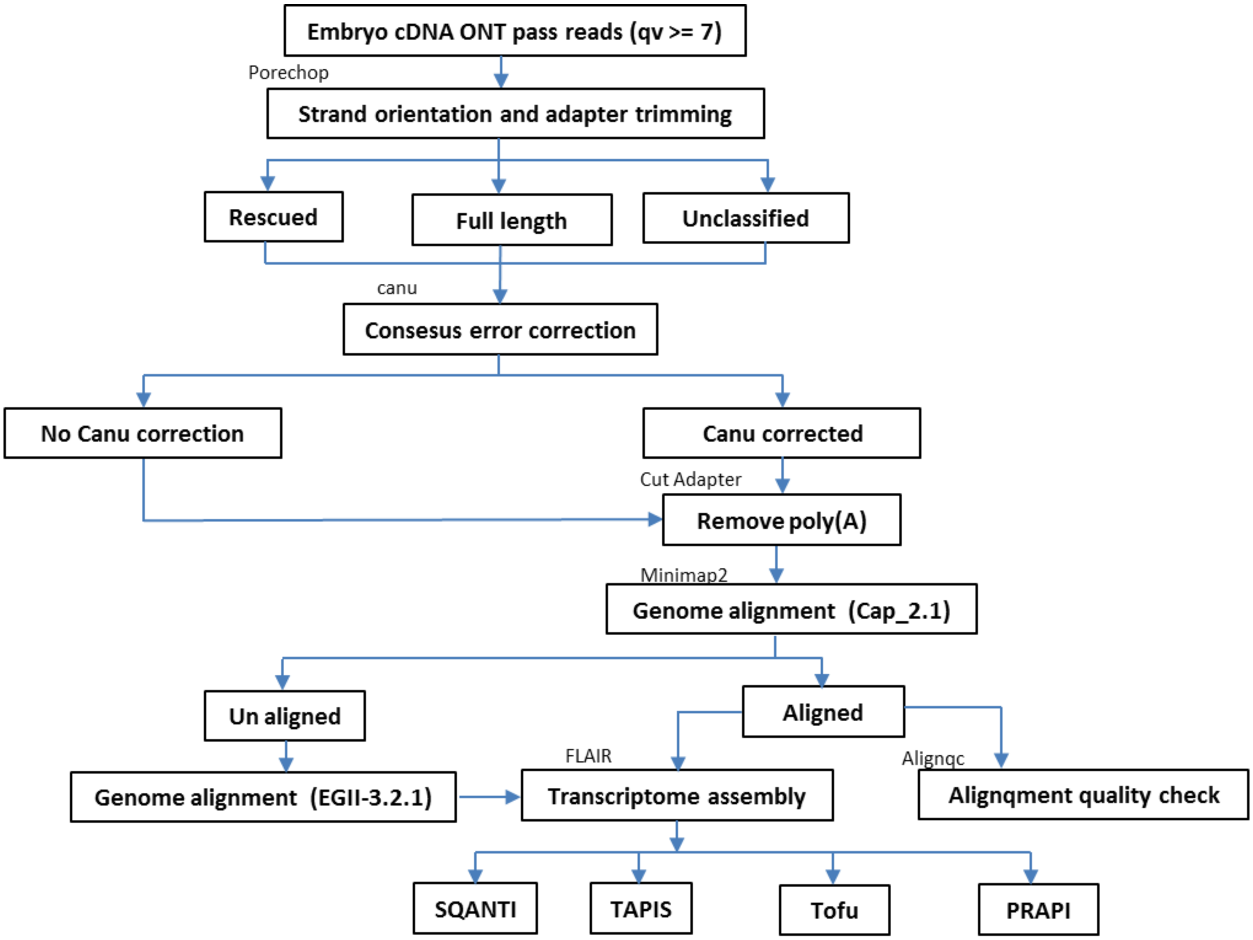


**Supplementary Figure S14: Data analysis workflow.** Pass reads (read with quality value equal or above 7) were processed using Porechop to remove adapters and also put them in their original orientation. Porechop classifies reads as ‘Full length’ (if both 5 and 3 prime adapters are identified), ‘Rescued’ (if adapters are identified within the gene sequence), and ‘Unclassified’ (if adapters are not confidently identified at both ends). All Porechop processed reads were passed to Canu for consesus error correction. Both error-corrected and non-corrected reads were combined and processed using Cut Adapter to remove the poly(A) tail. Processed reads were aligned using Minimap2 to the Cap_2.1 assembly () and the aligned reads collapsed into a transcriptome assembly using Flair. The assembly was further processed using SQANTI, Tofu, to assess quality of isoforms. Reads that did not align to Cap_2.1 assembly were aligned to EGII-3.2.1 assembly and the aligned reads collapsed using Flair.

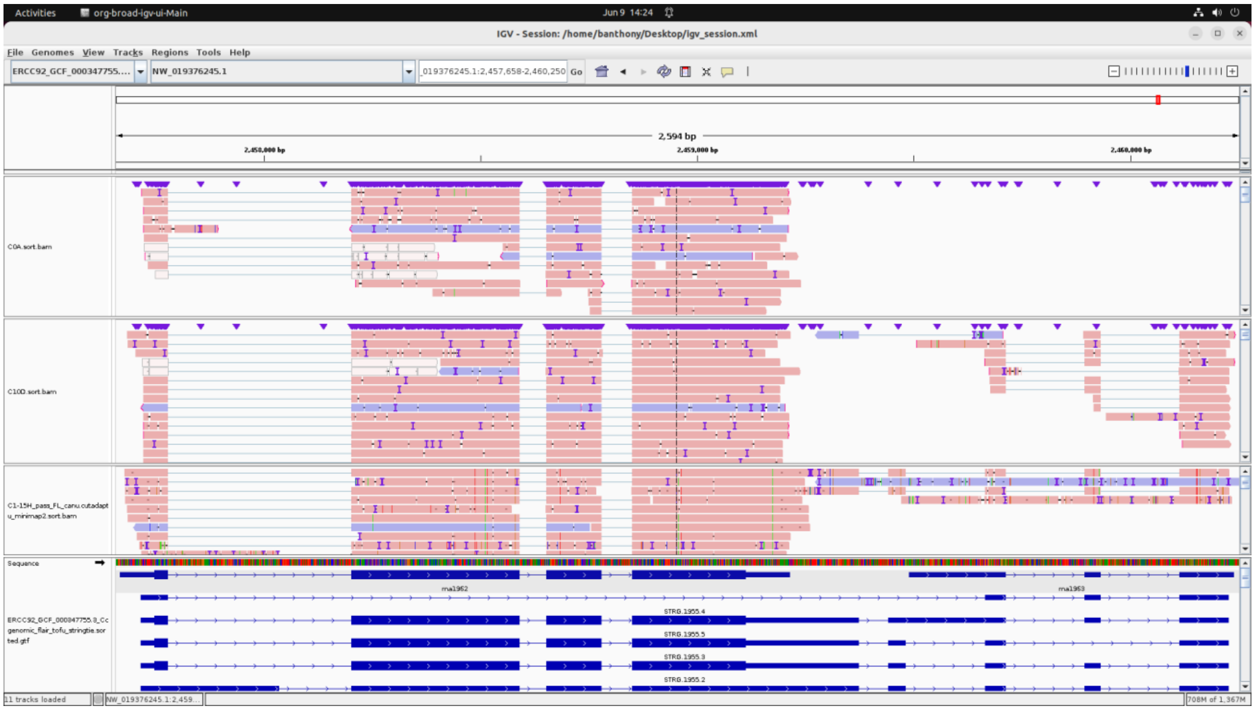

**Supplementary Figure S15:** Correction of NCBI gene models. We used SQANTI to identify NCBI gene models wrongly predicted to be separate genes but we could find a single read matching such genes. In this IGV screenshot, an example of such genes is shown.

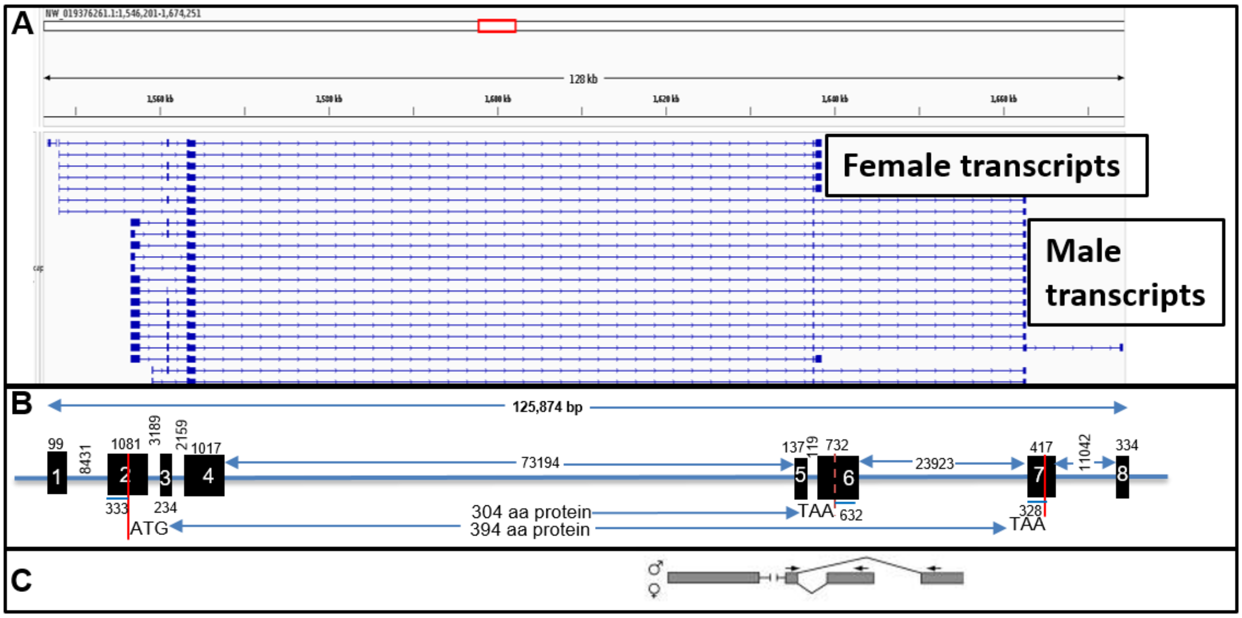


**Supplementary Figure S16: Gene model for Doublesex**

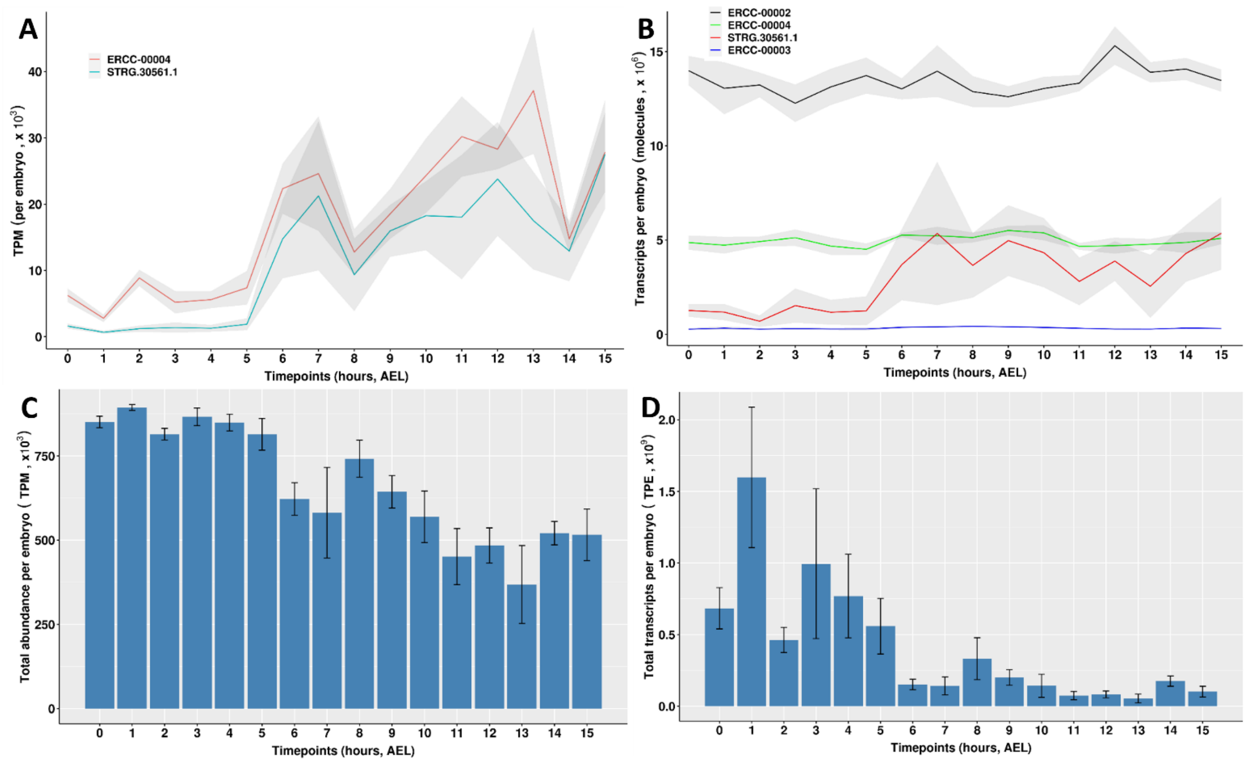


**Supplementary Figure S17: Comparison of relative and absolute gene expression quantification.** A) Relative normalization showing transcripts per million (TPM) across 16 time-points for one ERCC (ERCC-00004) and one transcript (STRG-30561.1). Mean (red line) and standard deviation (grey area) across the number of embryos used at each time-point are shown. B) Same as 'A' but showing absolute number of transcripts per embryo (TPE) for 3 ERCCs (ERCC-00002, ERCC-00004, ERCC-00003) and one transcript (STRG-30561.1). C) Bar graph showing the mean summed relative expression of each transcript (TPM) for each embryo across 16 time-points. Whiskers show standard deviation across the number of embryos at each time-point. D) Same as C but showing mean absolute number of transcripts per embryo.

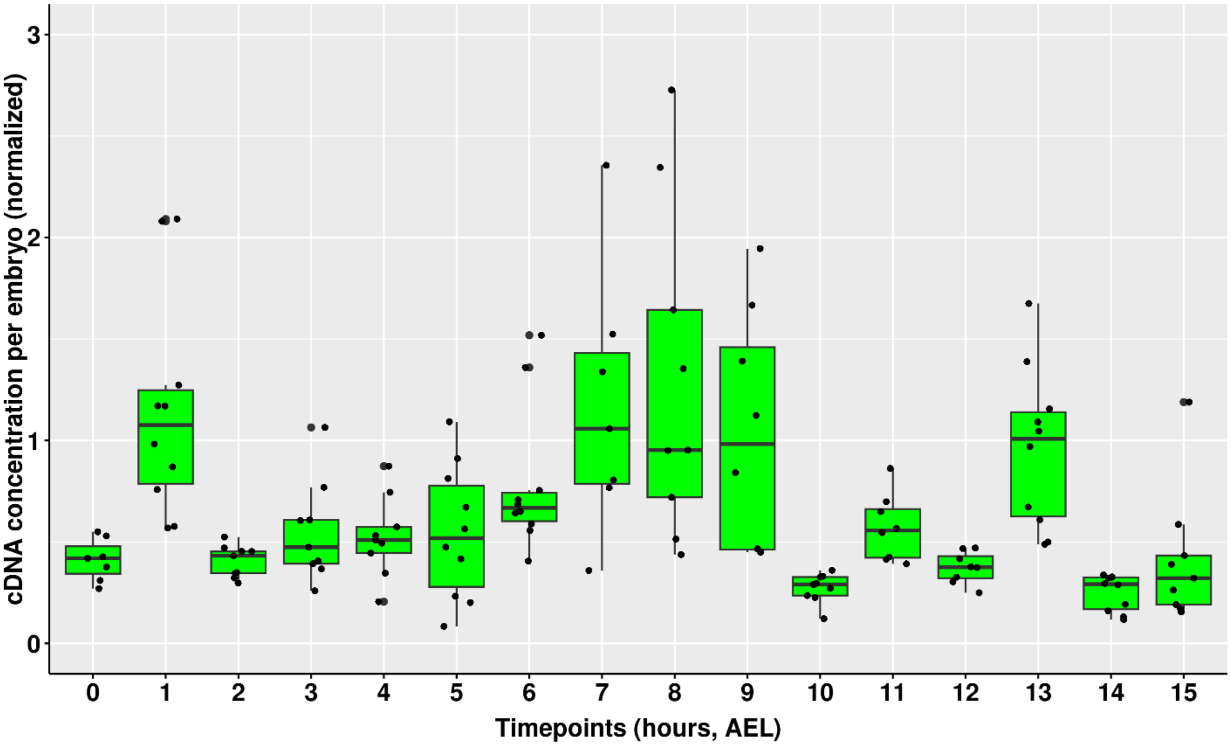


**Supplementary Figure S18:** Normalized cDNA amplicon yield. We prepared cDNA libraries from 12 embryos at each of the 16 time-points following our published protocol[13]. The concentration of libraries was estimated using Qubit dsDNA HS Assay kit (Thermo Fischer Scientific). Here, we show the concentration of each sample normalized for the total RNA extracted.

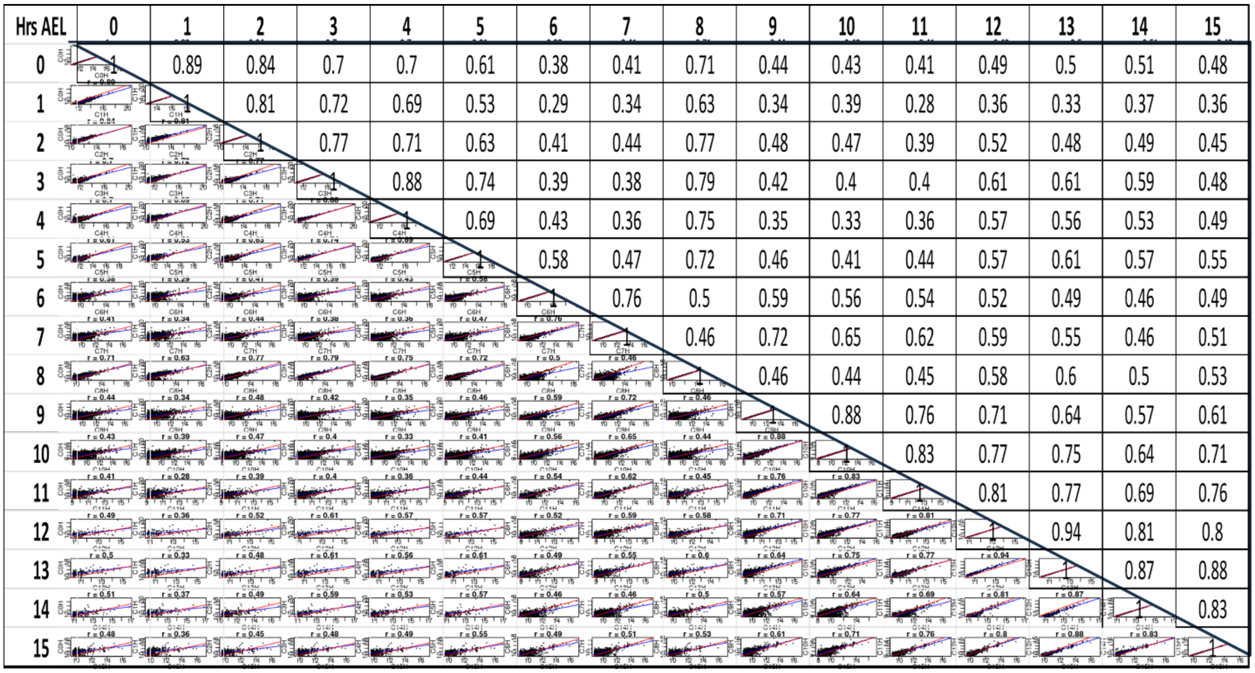


**Supplementary Figure S19:** Correlation of gene expression across time-points. The average isoform expression for each time-point was determined and used to calculate the Spearman correlation with the rest of the time-points. A hitman and raw correlation coefficients are shown.

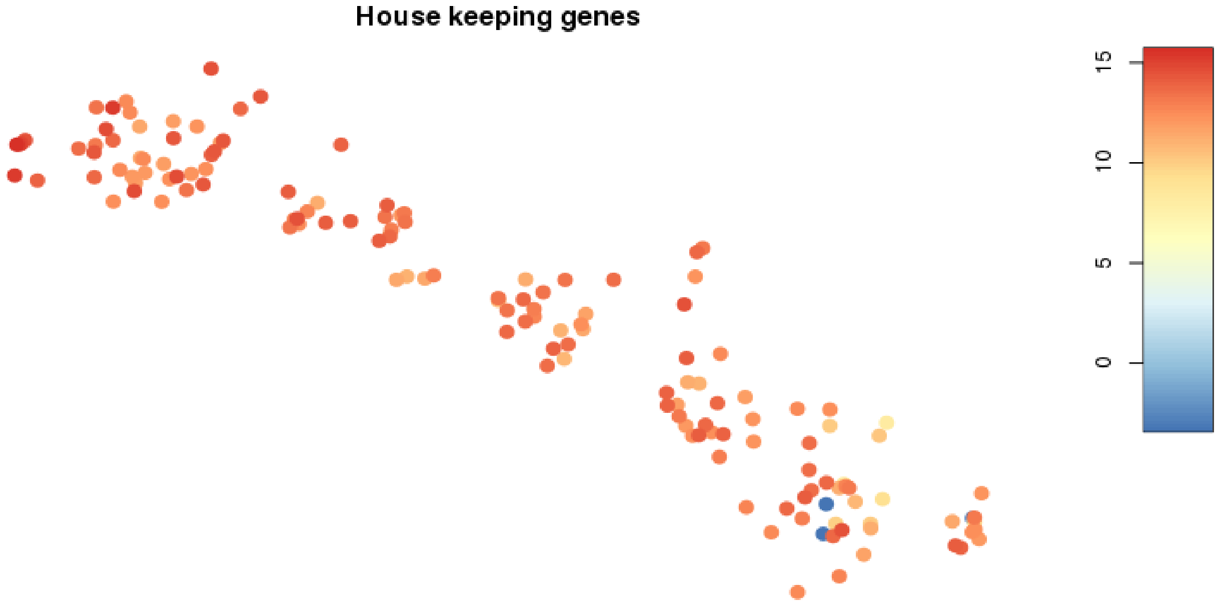


**Supplementary Figure S20: Quality control of the embryos.** We probed the expression of genes known to be involved in general cellular process including tubulin, RpL19, and RpL32.

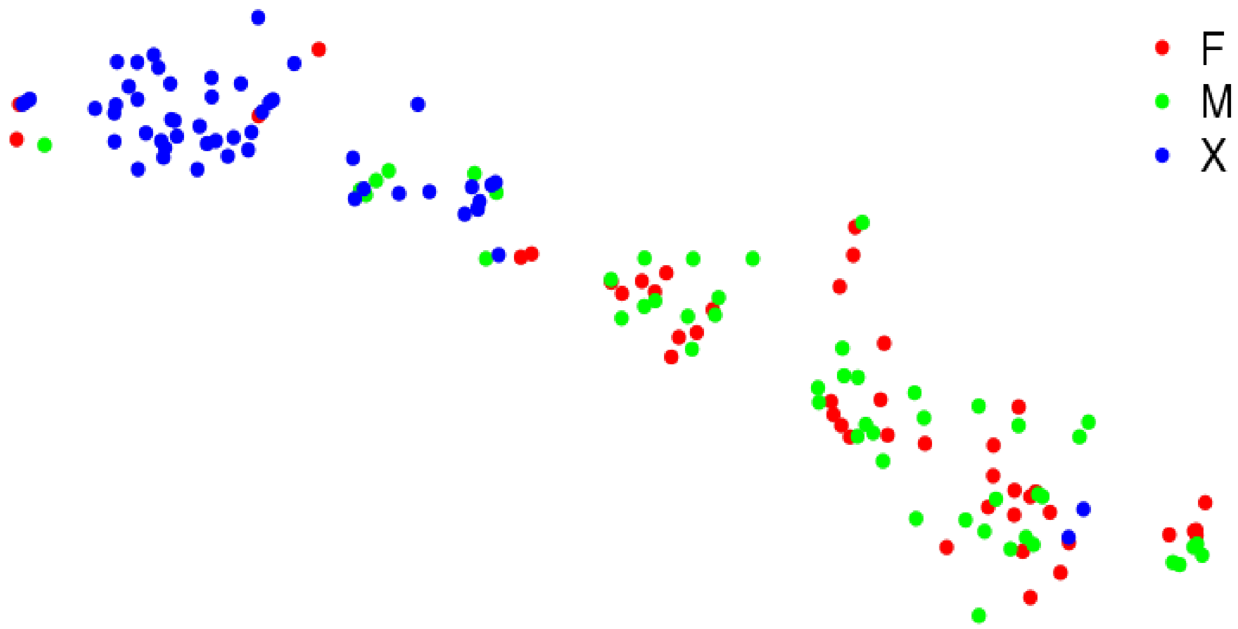


**Supplementary Figure S21: Sex assignment.** Embryos were sexed using genomic DNA extracted from each embryo and PCR of 2 genes, CcY and Y114. Following unsupervised clustering of the embryos based on their transcription profile, we called the sex of each embryo: red for female (F), green for male (M), and blue for undetermined sex (X).

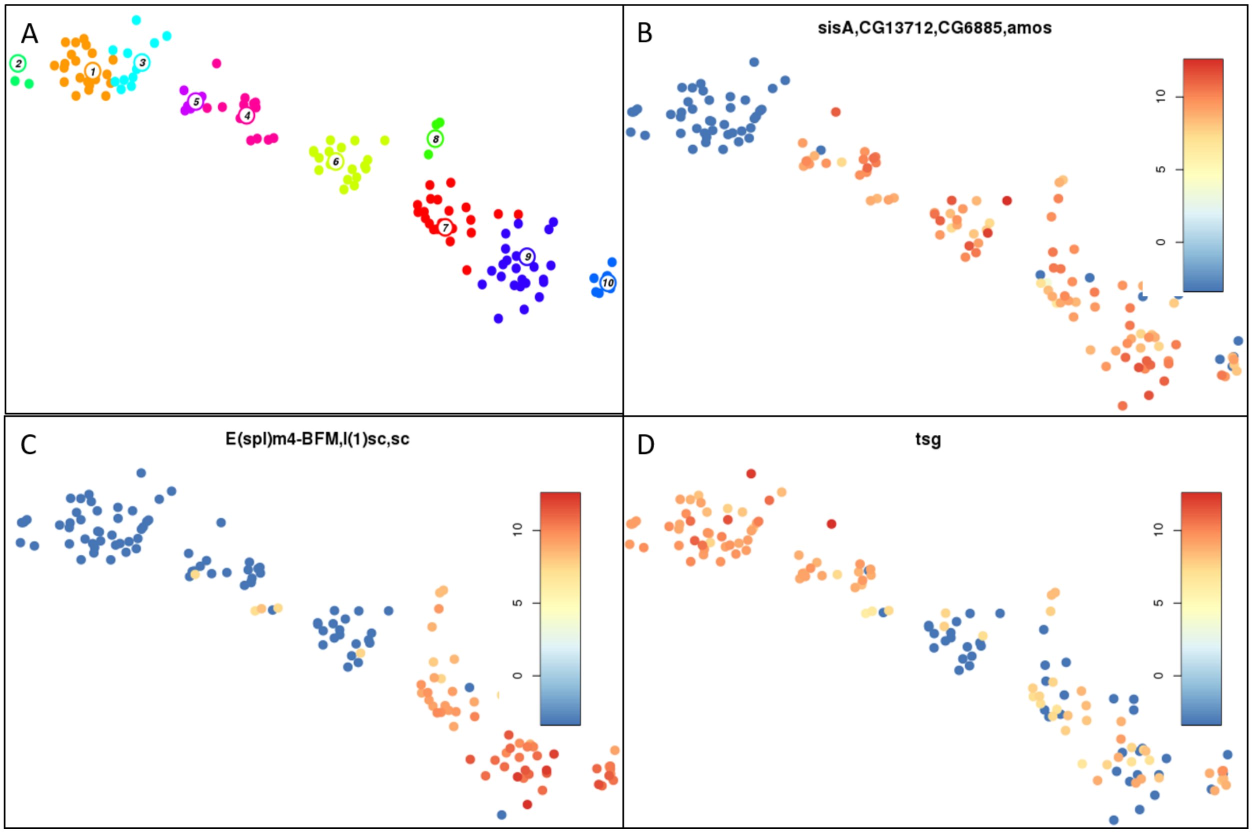


**Supplementary Figure S22: Probing genes involved in zygotic genome activation** [19, 20]. A) Unsupervised clustering of embryos as explained in Figure 2.. B) Identification of first expression of Drosophila minor wave of zygotic genome activation genes in medfly embryos. C) Identification of first expression of Drosophila major wave of zygotic genome activation genes in medfly embryos. D) Expression of tsg in medfly embryos.

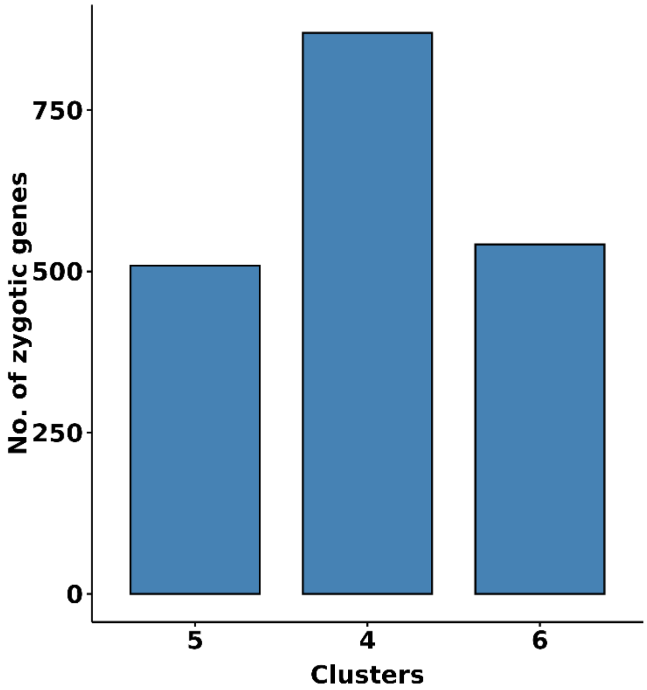


**Supplementary Figure S23: New zygotic genome genes.** We identified genes potentially transcribed from the zygotic genome. For all genes in clusters 5, 4, and 6, we determine genes that had not been previously detected in clusters 1, 2, or 3.

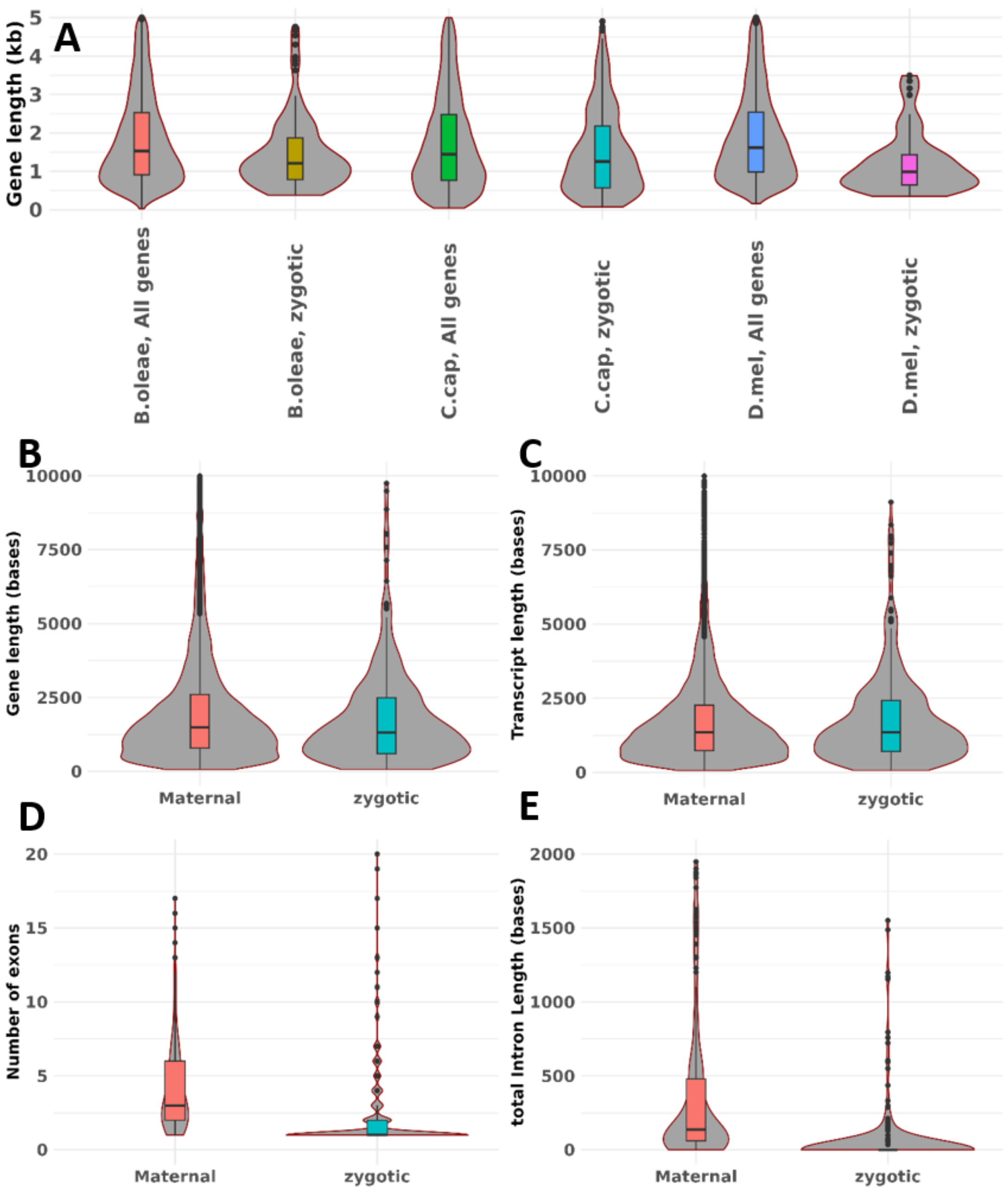


**Supplementary Figure S24: Zygotic gene length analysis**

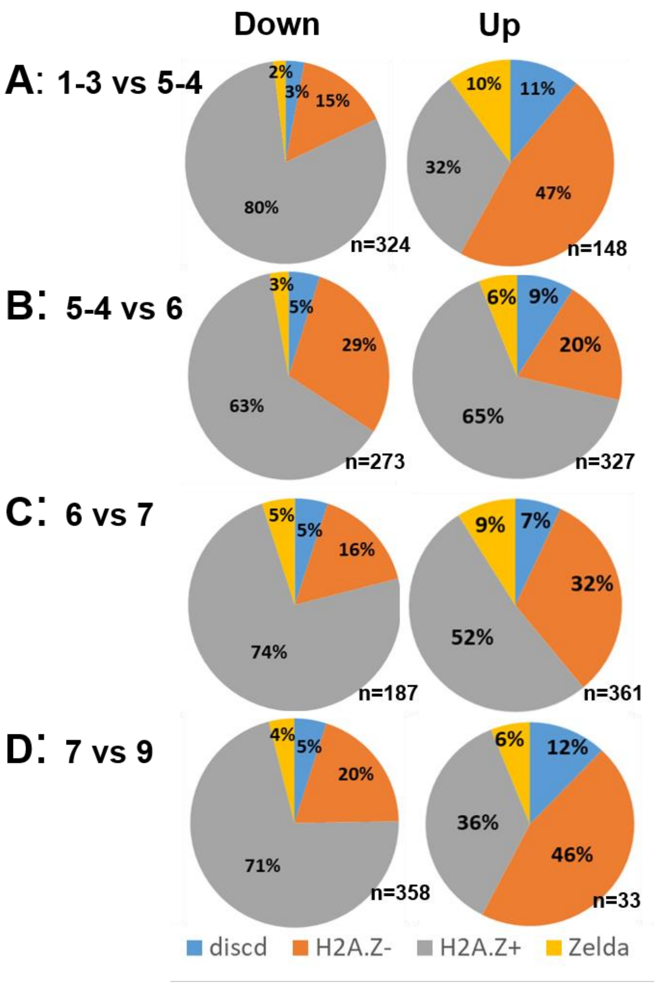


**Supplementary Figure S25: Probing of mechanisms for zygotic genome activation.** A-D) Differential gene expression was performed in successive clusters shown in Figure 2A. We combined clusters 1 and 3, and 5 and 4. We then determined significantly differentially expressed genes in clusters 1-3 versus 5-4 (A), 5-4 versus 6 (B), 6 versus 7 (C), and 7 versus 9 (D). We obtained the significantly differentially expressed genes and their *D. melanogaster* homologs. We annotated the homologs as to whether they contain Zelda motif or are enriched with H2A.Z or not (H2A.Z+ and H2A,Z-, respectively), or have discordant results [21, 22]. For each of the significantly down or upregulated genes in the clusters, we assigned them to their corresponding *D. melanogaster* annotation; Zelda motif, H2A.Z+, H2A.Z-, or discordant and determined the percentages.

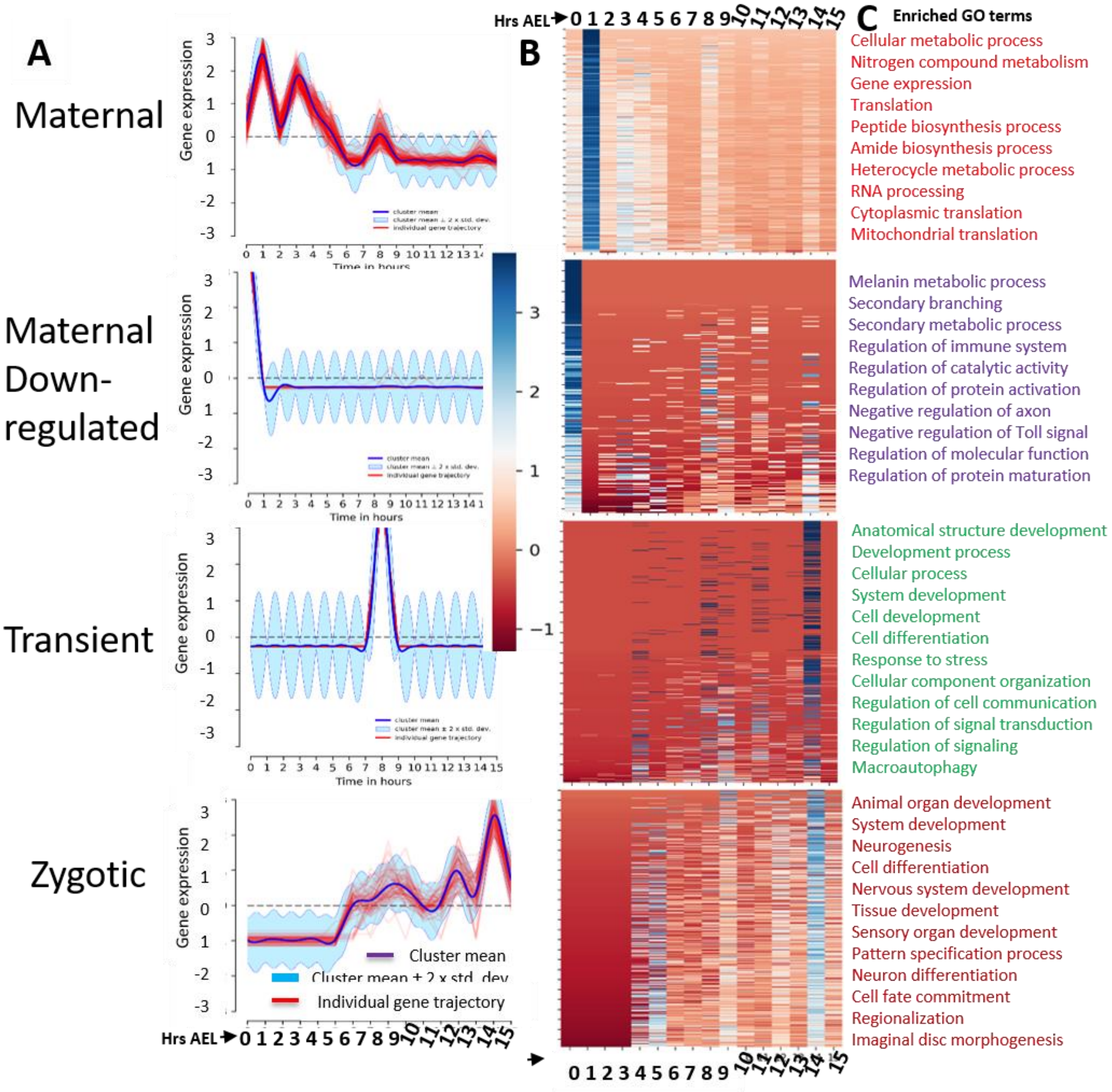


Supplementary Figure S26**:** Temporal clustering of gene expression**. A) The four temporal groups of clustering derived using real-time (as opposed to pseudo time) staging of embryo development. We used Drichlet process Gaussian process clustering**[68] **(DPGP) to put genes into clusters based on similarity in their temporal gene expression. We obtain 200 such clusters. We then further assigned these clusters to either of four groups: Maternal, Maternal Down-regulated, Transient, and Zygotic. A representative cluster is shown for each group.**

**B) Heatmap of gene expression. We combined all genes in each cluster and created a heatmap. C) Gene ontology. For each group. We used gProfiler to identify enriched biological processes with 5% false discovery rate, the top ones of which are shown. GO: gene ontology, Hrs: Hours, AEL: after egg laying.**

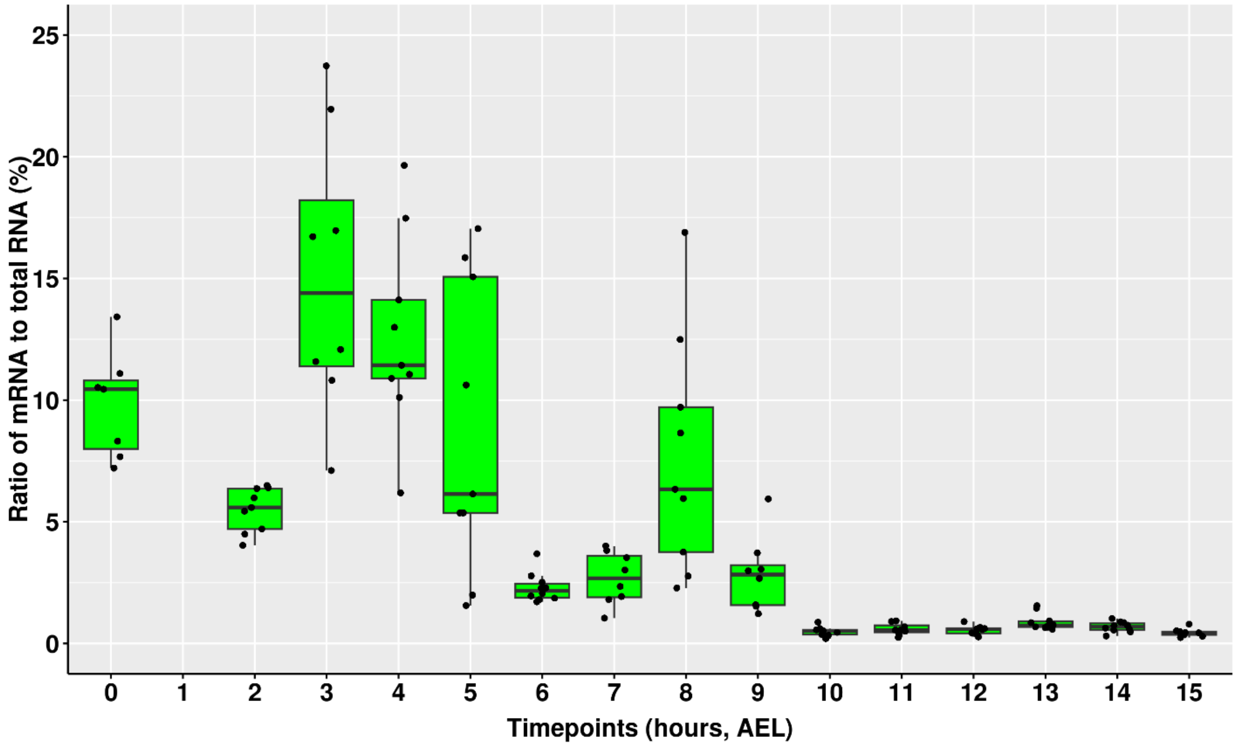


**Supplementary Figure S27:** Ratio of mRNA to total RNA. The computed total mRNA per embryo at each time-point was divided by empirically measured total RNA yield per embryo and shown here as a percentage.

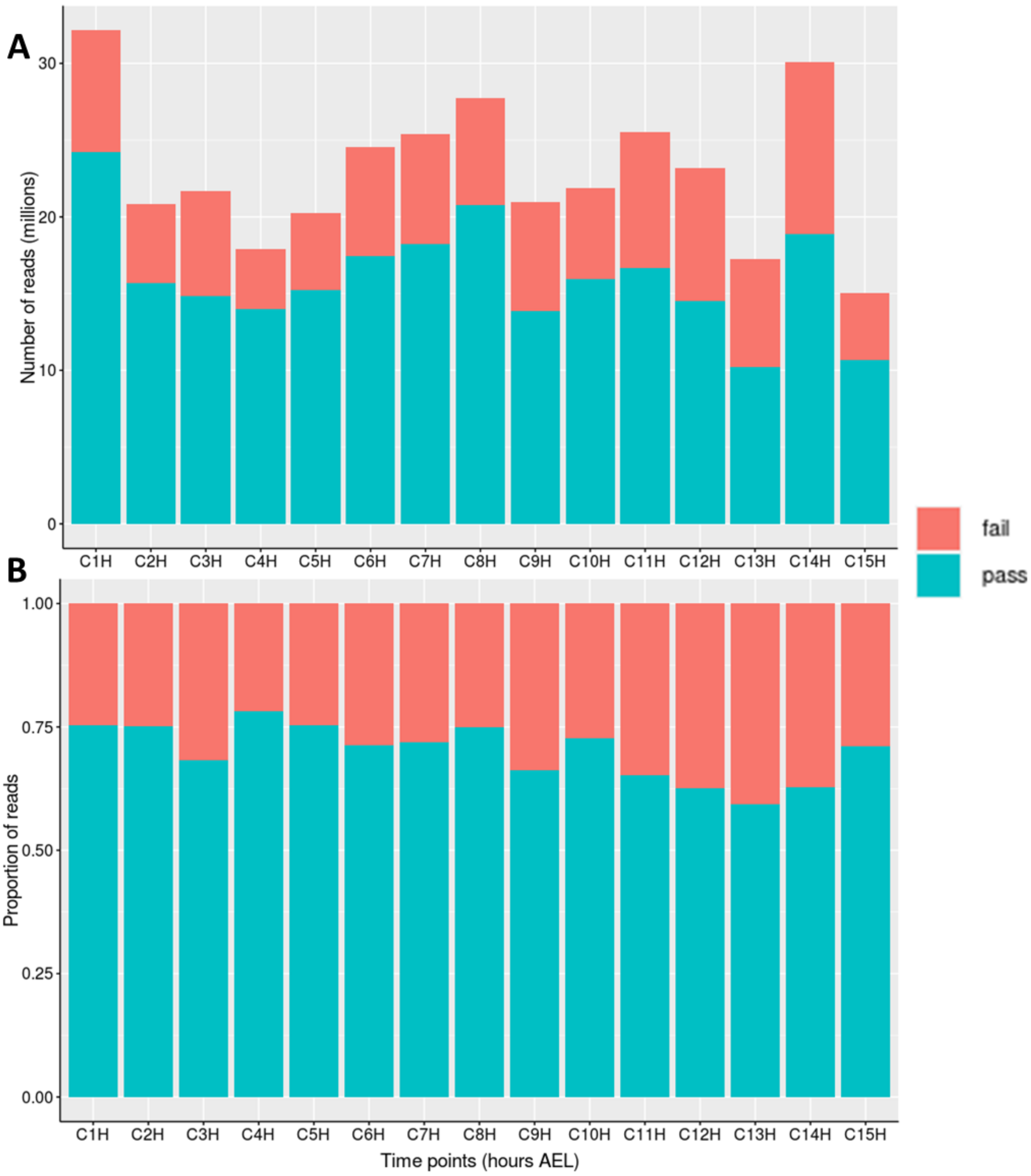


**Supplementary Figure S28: Total number of reads obtained from each time-point.** A) Total number of pass (reads with quality value, QV, of 7 and above) and fail reads (QV of less than 7) per time-point. B) Same as 'A' but showing proportion.

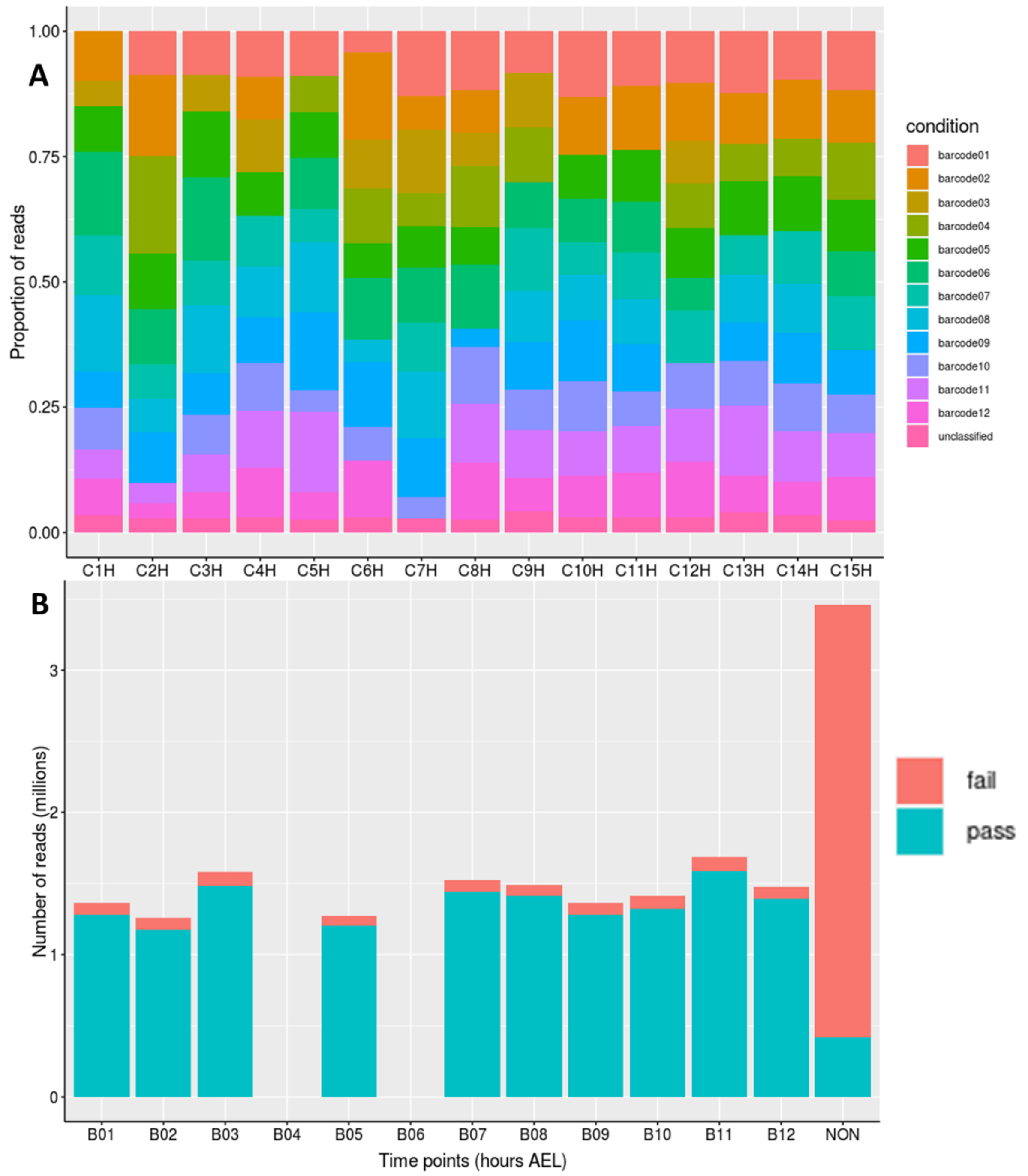


**Supplementary Figure S29: Barcode distribution of reads.** A) Proportional distribution of reads to the different barcodes across time-points. Unclassified refers to reads whose barcodes could not be reliably identified. Since each embryo was barcoded during PCR amplification of cDNA, we demultiplexed the samples. B) Distribution of reads to the different barcodes for the 4-hour time-point. Pass and fail reads are shown. The other time-points were not very different from this one which is shown as an example.

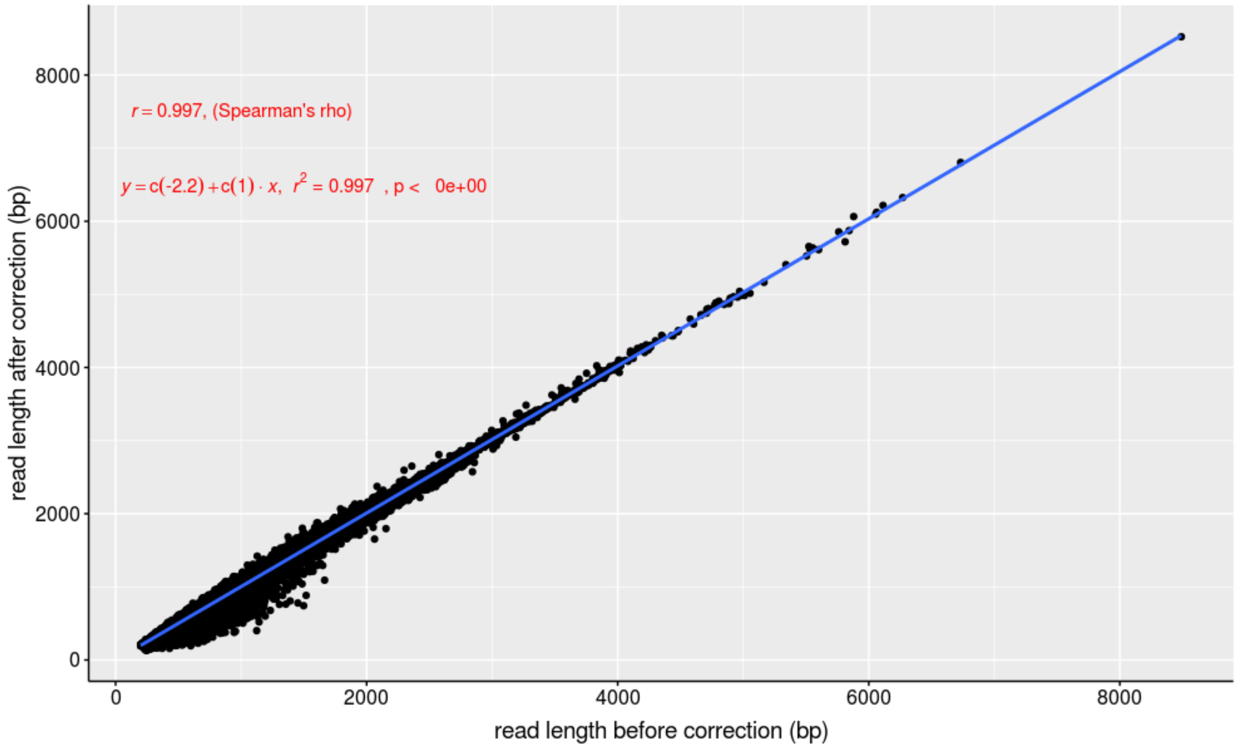


**Supplementary Figure S30: Correlation of read lengths before and after Canu correction.** All reads were supplied to Canu to perform consensus error correction. The lengths of reads that were corrected were compare before and after correction and results are shown.

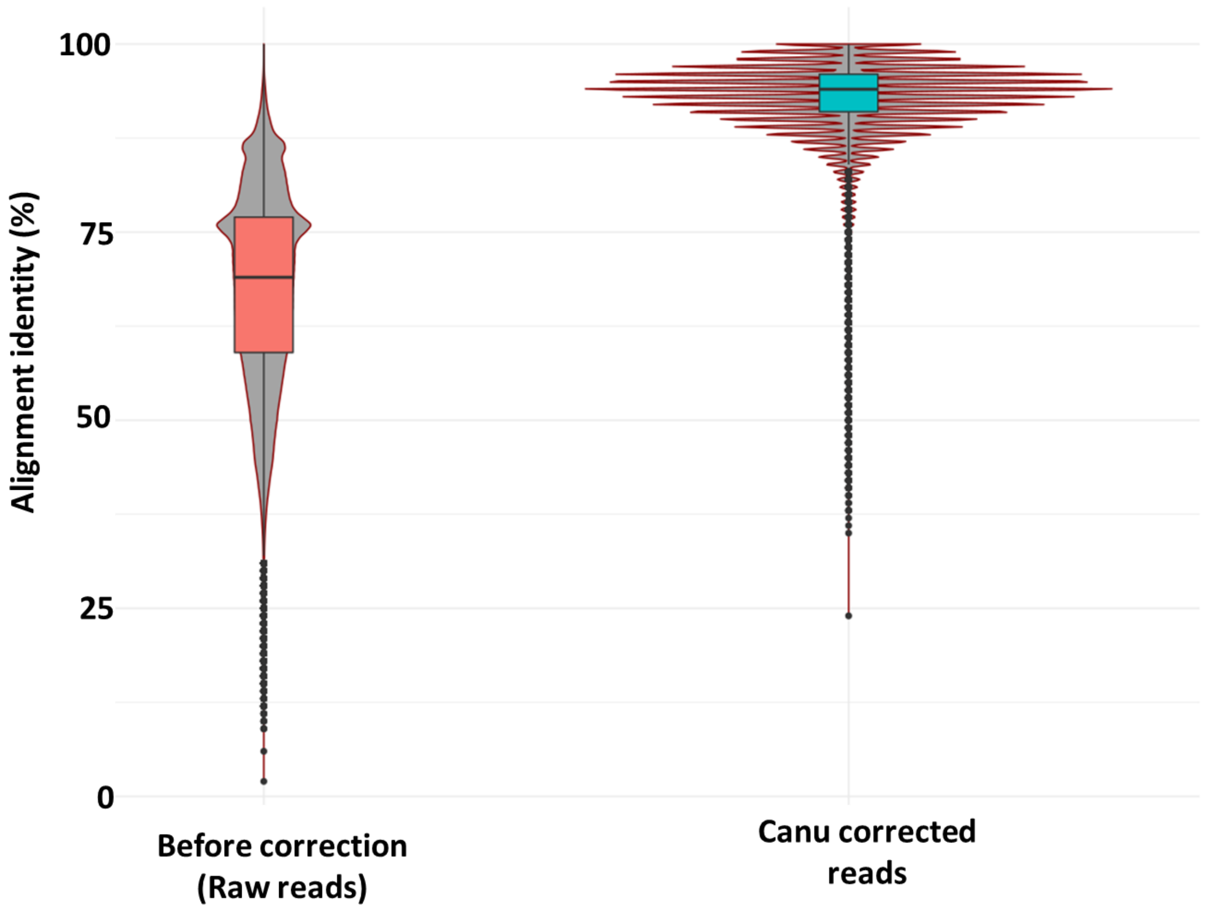


**Supplementary Figure S31: Alignment identity of reads before and after Canu correction.** Reads were supplied to Canu for consensus error correction. The alignment identity of error corrected reads to the Medfly Cap_2.1 genome before and after error correction was determined and shown here.

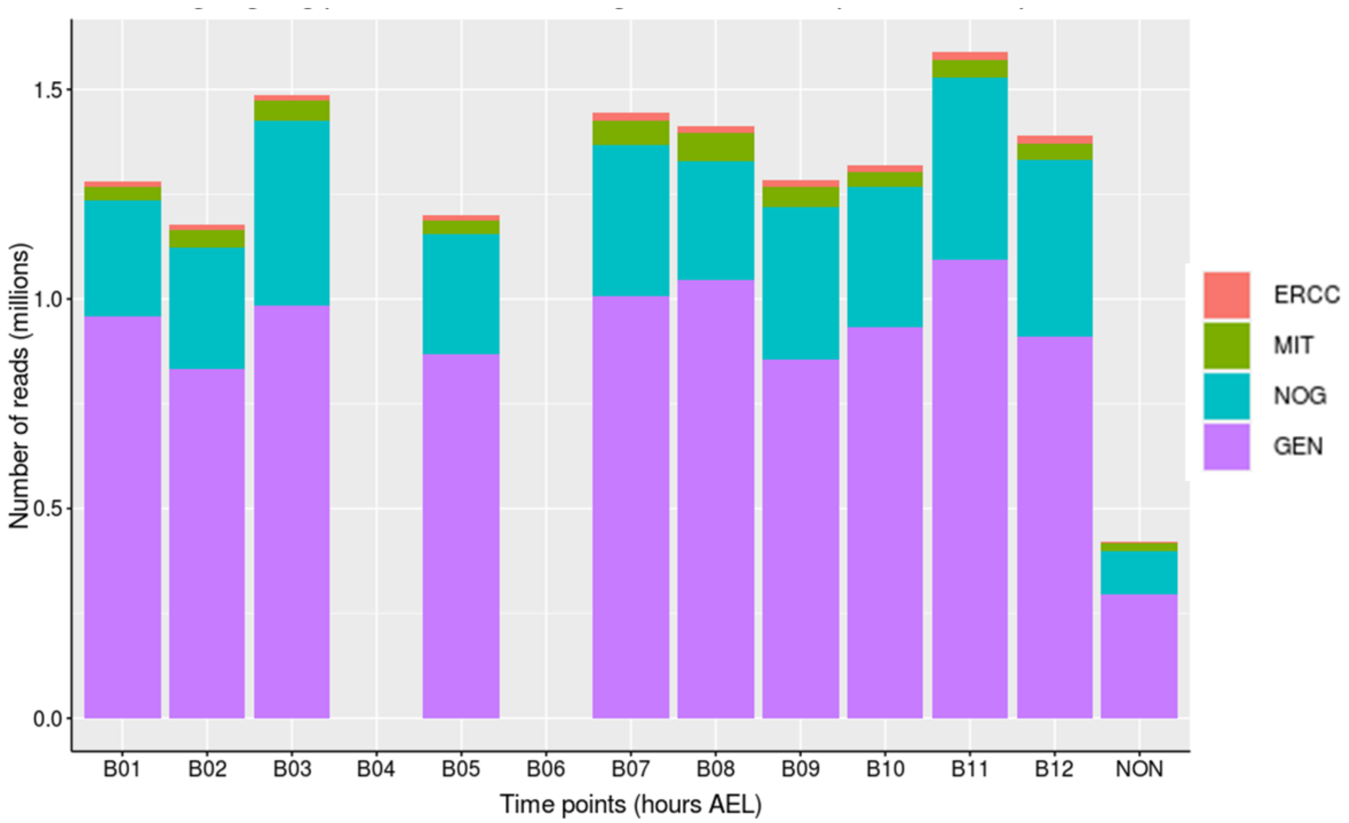


**Supplementary Figure S32: Distribution of reads across genomic features.** Reads were aligned to the genome assembly supplemented with ERCC external quality sequences. Reads that aligned to each of the 4 genomic features; ERCC, mitochondrial genome (MIT), genome (GEN) and unaligned reads (NOG) were counted. This figure shows an example for one of the time-points.

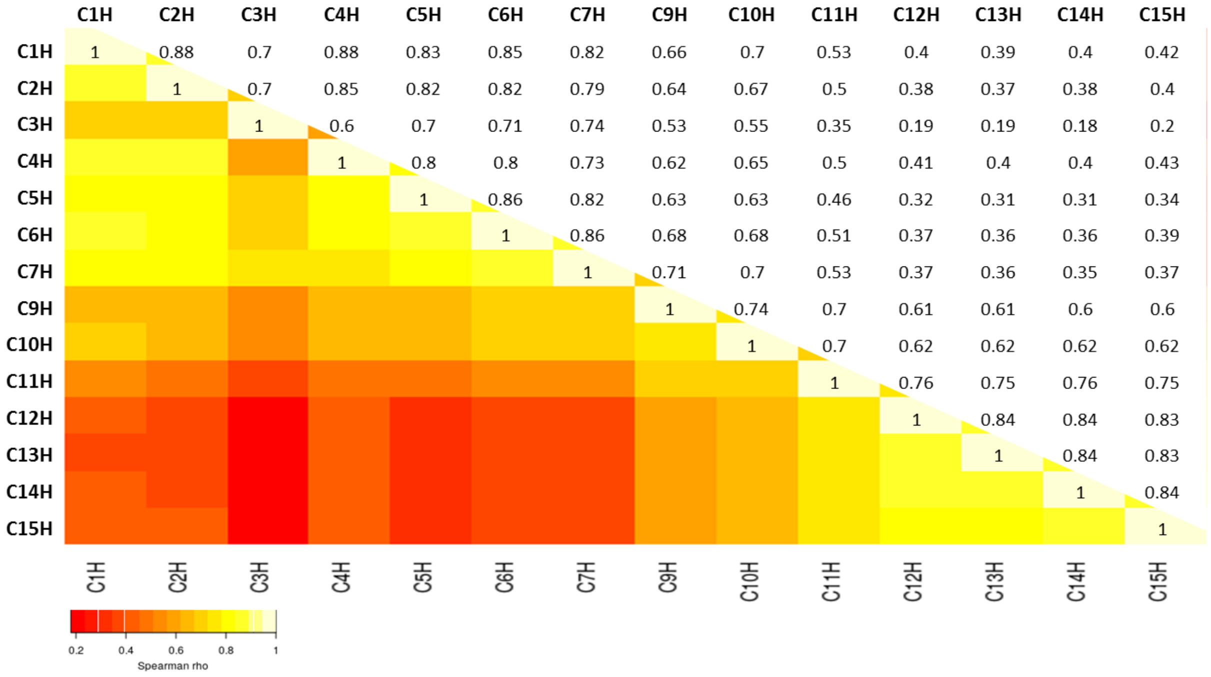


**Supplementary Figure S33: Correlation of gene expression across time-points.** The average isoform expression for each time-point was determined and used to calculate the Spearman correlation with the rest of the time-points. A hitman and raw correlation coefficients are shown.

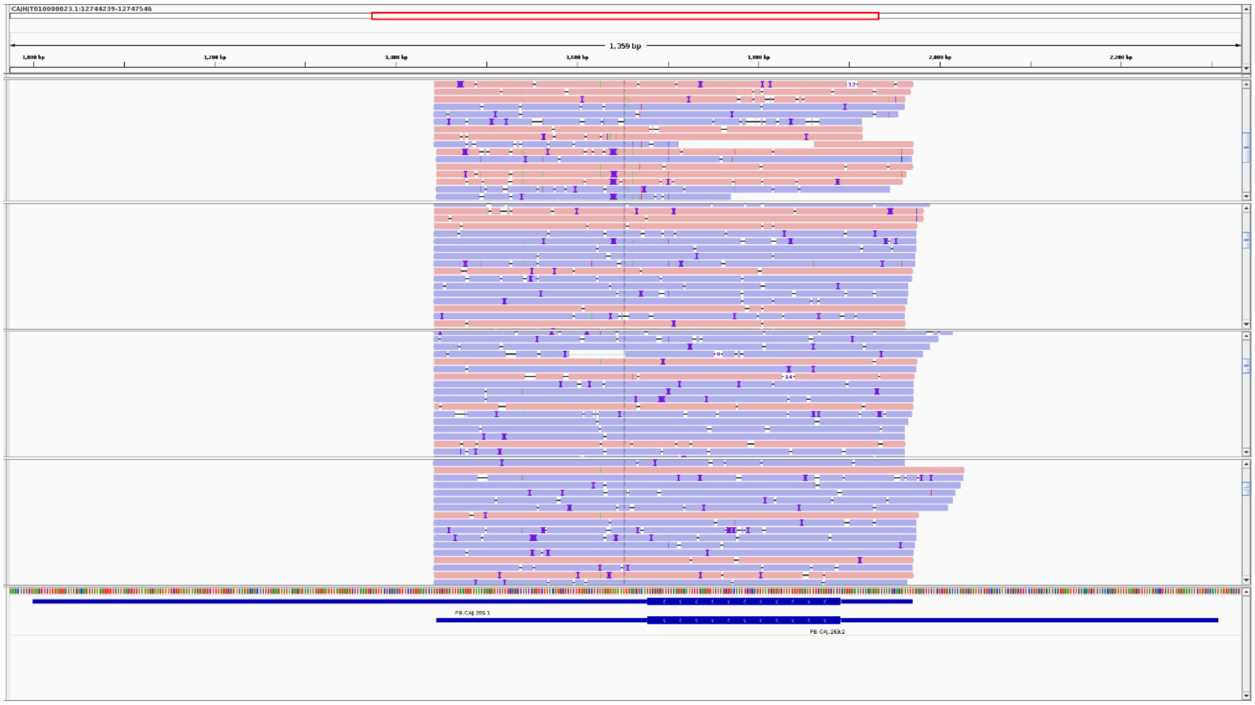

**Supplementary Figure S34: *Ceratitis capitata* maleness-on-the Y (CcMoY) transcripts.** Reads obtained from embryos collected at 4 hours after egg laying (AEL) were aligned to the CcMoY gene identified in the EGII-3.2.1 genome. Integrative genome viewer (IGV) is shown here with 4 tracks corresponding to 4 different single male embryos. Reads were not oriented prior to alignment.

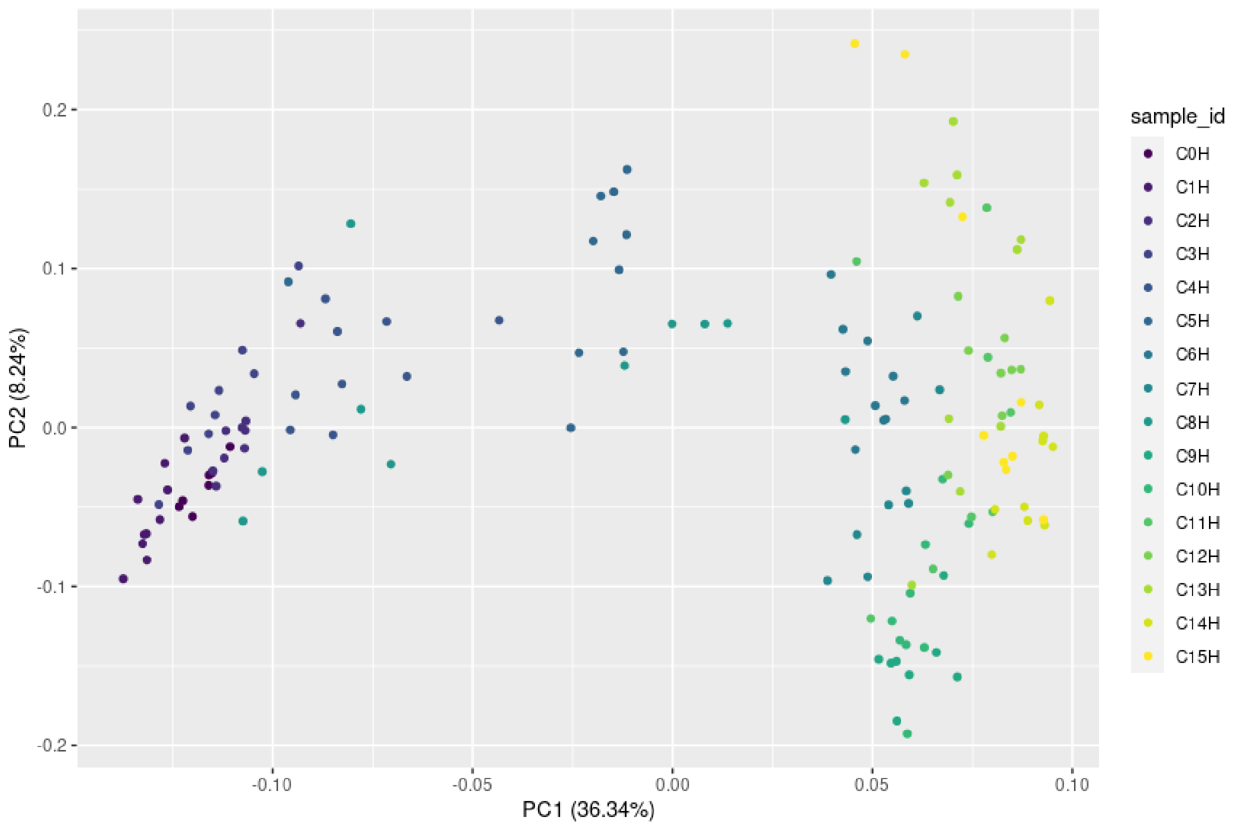


**Principal component analysis (PCA) of average gene expression (transcripts per embryo) at each time-point**
